# Neural correlates of perceived emotions in human insula and amygdala

**DOI:** 10.1101/2022.01.29.477631

**Authors:** Yang Zhang, Wenjing Zhou, Juan Huang, Bo Hong, Xiaoqin Wang

## Abstract

The emotional status of a speaker is an important non-linguistic cue carried by human voice and can be perceived by a listener in vocal communication. Understanding the neural circuits involved in processing emotions carried by human voice is crucial for understanding the neural basis of social interaction. Previous studies have shown that human insula and amygdala responded more selectively to emotional sounds than non-emotional sounds. However, it is not clear whether the neural selectivity to emotional sounds in these brain structures is determined by the emotion presented by a speaker or by the emotion perceived by a listener. In this study, we recorded intracranial electroencephalography (iEEG) responses to emotional human voices while subjects performed emotion recognition tasks. We found that the iEEG responses of Heschl’s gyrus (HG) and posterior insula were determined by the presented emotion, whereas the iEEG responses of anterior insula and amygdala were driven by the perceived emotion. These results suggest that the anterior insula and amygdala play a crucial role in representing subjectively perceived emotion.

## Introduction

Human voice contains not only communicative information, but also information for recognizing individual identities and their emotional status (Ellis and Andrew, 1989). The ability to perceive the emotional information contained in human voice is critical for social interactions. Previous studies have shown that human amygdala plays an important role in recognizing emotions from facial expressions (Adolphs, 2010; Wang et al., 2014; Morris et al., 1998). Earlier studies suggested that the human amygdala show stronger responses to fearful faces than to faces with other emotional expressions (Morris et al., 1996); while recent studies supported the idea that the human amygdala responds to facial expressions of all types of emotions (Fitzgerald et al., 2006). In addition, single neurons’ responses in the human amygdala were found to encode subjective perception of emotional faces (Wang et al., 2014; Morris et al., 1999; Sander and Scheich, 2001). Furthermore, neuroimaging studies have also suggested the important role of amygdala in perceiving vocally expressed emotions (Phillips et al., 1998; Morris et al., 1999; Sander and Scheich, 2001), in which enhanced responses to emotional voices were observed compared with neutral voices. Therefore, there is substantial evidence suggesting the involvement of human amygdala in emotion processing.

In addition to the human amygdala, the human insula also plays an important role in emotion processing. The human insula lies under the frontal, parietal, and temporal opercula. Studies have revealed that human insula is involved in a wide range of tasks and conditions, including sensory representations (Ostrowsky et al., 2002; Remedios et al., 2009; Augustine, 1996; Bamiou et al., 2003), affective representations (Zaki et al., 2012; Phan et al., 2002), and other high-level cognitive functions (Menon and Uddin, 2010; Sridharan et al., 2008). Tracing studies in non-human primates have revealed extensive reciprocal connections between the auditory cortex (Augustine, 1985; Mesulam and Mufson, 1985) and the insula, as well as between the insula and the amygdala (Mufson et al., 1981; Augustine, 1985; Höistad and Barbas, 2008). A recent human study in our lab have learned that the posterior insula (PI) is structurally and functionally connected with the auditory cortex, whereas the anterior insula (AI) is connected with the amygdala. Comparing with non-emotional sounds, emotional sounds induced enhanced neural responses in the anterior insula and the amygdala (Zhang et al., 2019), suggesting that the anterior insula and the amygdala in human are involved in auditory emotion processing. However, it remains unknown whether the emotion-selective responses are driven by the emotion presented by the speaker or perceived by the listener. It has been suggested that neurons in Heschl’s gyrus (HG, human primary auditory cortex) encode objective acoustic features of sounds (Chang et al., 2010; Mesgarani et al., 2014; Brugge et al., 2009; Griffiths et al., 2010); whereas neurons in amygdala encode subjective perception of emotional faces (Wang et al., 2014; Morris et al., 1999; Sander and Scheich, 2001). Given the different connectivity patterns of anterior and posterior insula (AI connects to amygdala; PI connects to HG), we hypothesized that the anterior insula and amygdala encode the emotion perceived by the listener, whereas the posterior insula and HG encode the emotion presented by the speaker which depends on the acoustic features of the auditory stimuli.

To test the hypothesis, we combined intracranial electroencephalography (iEEG) recordings from epilepsy patients implanted with depth electrodes in the insula (PI and AI regions), amygdala, and HG (Figures 1A, B) with their behavior performances in auditory emotion recognition tasks. The aim of the present study is to differentiate the objective acoustic processing roles (representation of the presented emotion of the stimuli that depend on acoustic features) and the subjective emotion perception roles (representation of the perceived emotion) of several brain areas (HG, PI, AI, and amygdala) for auditory emotion recognition. iEEG signals were recorded from 24 patients while emotional human voices and morphed emotional human voices were delivered to them. They were asked to tell which types of auditory emotion they perceived following each stimulus being presented. By comparing iEEG signals corresponding to correct emotion categorizations trials (the perceived emotion by the listener is the same as the presented emotion by the speaker) with those to perception errors (the perceived emotion by the listener is not the same as the presented emotion by the speaker), we were able to dissociate the electrophysiological markers of the acoustic processing of emotional voices (presented emotion by the speaker) and the perception of emotion (perceived emotion by the listener). We showed that responses in the posterior insula and HG are more associated with the emotion presented by the speaker, whereas responses in the anterior insula and amygdala depend more on the perception of emotion by the listener. Our findings suggest a role of human insula and amygdala in representing the subjective perception of auditory emotion.

**Figure 1.**
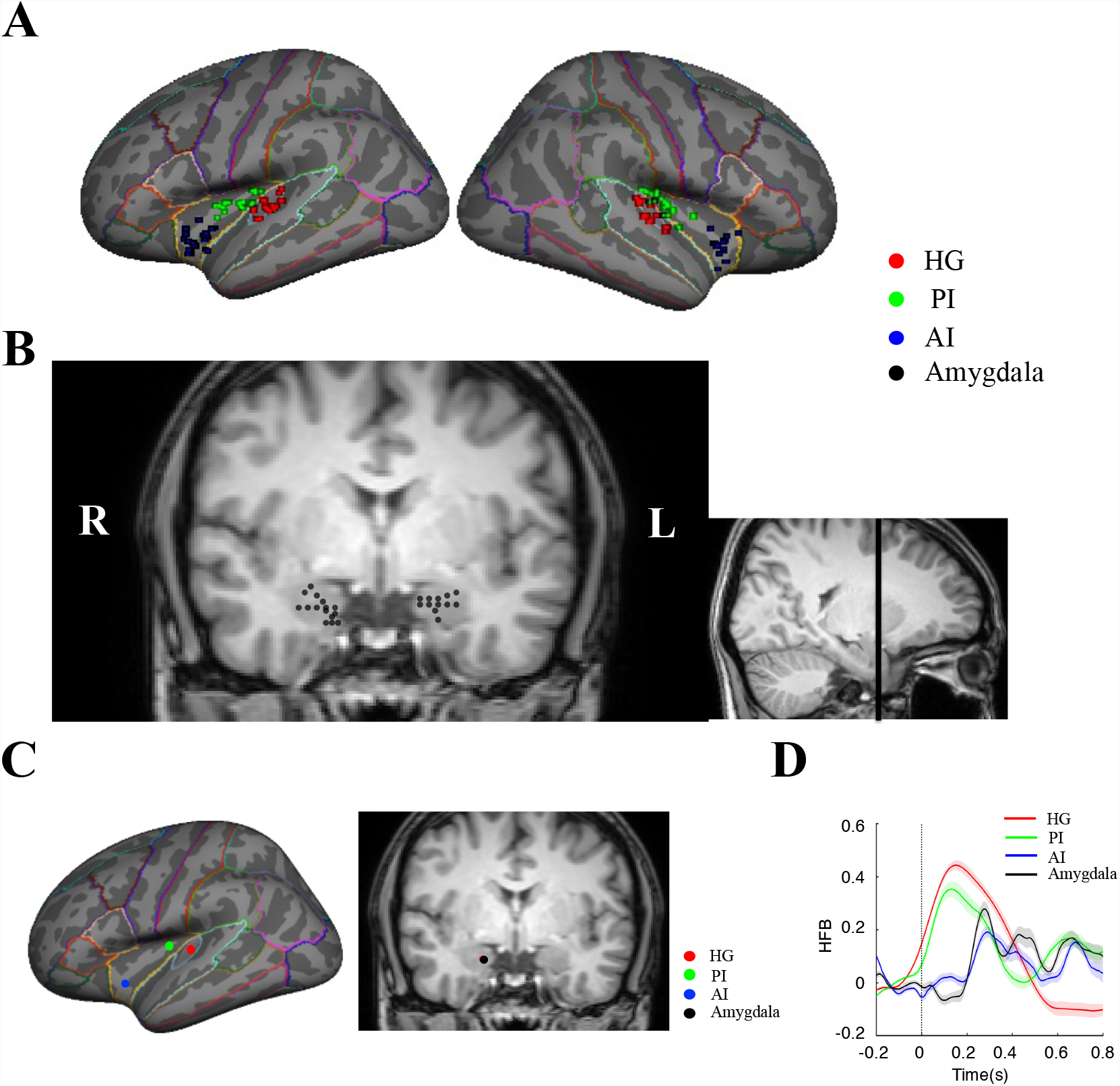
Locations of all recorded electrodes and responses of single electrodes in HG, PI, AI, and amygdala: (A) Locations of electrodes in HG, PI, and AI on inflated brain surface (total electrode number: HG (28), PI (30), AI (26)). (B) Locations of electrodes in amygdala (total electrode number: Amygdala (27)). (C) Example electrodes locations in HG (red dot), PI (green dot), AI (blue dot), and amygdala (black dot). (D) Time series of the HFB responses of the example electrodes in (C) to an emotional stimulus (“disgust (2)”; mean ± sem (standard error of mean)).

## Materials and Methods

### Participants

24 patients (mean age: 24.5, 13 males and 11 females) with intractable epilepsy were recruited in the study. Each patient was implanted with intracranial depth iEEG electrodes (0.8 mm electrode diameter and 3.5 mm inter-electrode center-to-center distance) as part of their clinical treatments for epilepsy (see Table 1 for the electrode coverage of each patient). Two experiments were conducted in this study. All patients participated in Exp. 1, the Emotion Recognition Experiment, and 7 of them (mean age: 26.7, 3 males and 4 females) participated in Exp. 2, the Morphing Experiment (Table 1). Patients were all right-handed with normal Intelligence quotients and normal hearing. Written informed consent was obtained from each subject before enrollment. The experimental protocol was approved by the Institutional Review Board at Tsinghua University and the affiliated Yuquan Hospital.

**Table 1.** Clinical profiles of the epilepsy patient subjects who underwent iEEG recordings

| Subject | Age | Gender | Seizure Focus Location | Electrodes Coverage in insula, amygdala, and HG | Emotion Recognition Experiment | Morphing Experiment | Language Dominance |
| --- | --- | --- | --- | --- | --- | --- | --- |
| S1 | 38 | Female | Left hippocampus | Left amygdala, Left PI, Left HG | √ | × | Left |
| S2 | 28 | Male | Right hippocampus | Right amygdala, Right PI, Right HG | √ | × | Left |
| S3 | 30 | Female | Right temporal lobe | Right amygdala, Right PI, Right HG | √ | × | Left |
| S4 | 19 | Male | Left posterior temporal lobe | Left AI | √ | × | Left |
| S5 | 31 | Male | Left parietal lobe | Left amygdala, Left PI, Left HG | √ | √ | Left |
| S6 | 24 | Male | Left posterior temporal lobe | Left amygdala, Left PI, Left HG | √ | × | Left |
| S7 | 15 | Female | Right temporal lobe | Right PI, Right HG | √ | × | Left |
| S8 | 19 | Male | Left hippocampus | Left amygdala, Left AI | √ | × | Left |
| S9 | 19 | Female | Right posterior temporal lobe | Right PI, Right HG | √ | √ | Left |
| S10 | 15 | Male | Right frontal lobe | Right amygdala, Right AI | √ | × | Left |
| S11 | 24 | Female | Left hippocampus | Left amygdala, Left PI, Left HG | √ | × | Left |
| S12 | 21 | Male | Left hippocampus | Left amygdala, Left AI, Left PI, Left HG | √ | × | Left |
| S13 | 22 | Male | Left hippocampus | Left and right amygdala, Left and right PI | √ | × | Left |
| S14 | 24 | Female | Right hippocampus | Right PI, Right AI, Right HG | √ | × | Left |
| S15 | 47 | Female | Left anterior temporal lobe | Left amygdala, Left AI | √ | √ | Left |
| S16 | 16 | Female | Right occipital lobe | Right PI, Right HG | √ | √ | Left |
| S17 | 25 | Male | Right parietal lobe | Right PI, Right AI, Right HG | √ | × | Left |
| S18 | 13 | Female | Left occipital lobe | Left PI, Left HG | √ | × | Left |
| S19 | 26 | Male | Left hippocampus | Left amygdala, Left PI, Left HG | √ | × | Left |
| S20 | 27 | Male | Left hippocampus | Left and right amygdala | √ | × | Left |
| S21 | 33 | Female | Left parietal lobe | Left PI, Left HG | √ | × | Left |
| S22 | 33 | Male | Right hippocampus | Left and right amygdala, Left and right AI | √ | √ | Left |
| S23 | 25 | Female | Right premotor cortex | Right amygdala, Right PI, Right AI, Right HG | √ | √ | Left |
| S24 | 16 | Male | Right anterior temporal lobe | Right amygdala, Right PI, Right AI, Right HG | √ | √ | Left |

### iEEG data acquisition and electrode localization

iEEG signals were recorded via a 256-channel Neurofax EEG-1200c amplifier/digitizer system (Nihon Kohden, Japan) from implanted depth electrodes. The data were sampled at 2000Hz. Electrodes that located in the insula, HG, and amygdala were included in the analysis. The locations of the electrodes were determined using Freesurfer (http://surfer.nmr.mgh.harvard.edu/). We reconstructed the individual brain’s three-dimensional-map with electrode locations on the surface by aligning the pre-surgical high-resolution T1-weighted magnetic resonance imaging (MRI) obtained using a Philips Achieva 3.0T TX scanner with the post-surgical computed tomography (CT) images obtained using the Siemens SOMATOM Sensation 64 CT scanner. This registration was visually verified and manually adjusted if necessary (Ding et al., 2019; Zhang et al., 2021). Furthermore, we co-registered the individual MRIs to the fsaverage brain through Freesurfer to show all electrodes on an average brain surface. PI and AI were determined according to the resting state functional magnetic resonance imaging (rsfMRI) data of these subjects, as previously described (Zhang et al., 2019). HG, insula, and amygdala were defined and dissociated according to the anatomic labels of the average brain in Freesurfer.

### Experimental design and statistical analysis

#### Stimuli and Task

The stimuli consisted emotional voices that were chosen from the Montreal Affective Voices (Belin et al., 2008), in which actors or actresses were instructed to produce the vowel /a/ with different emotional interjections. In Exp. 1, the emotion recognition experiment, stimuli were 12 human voices produced by two identities (one male and one female) expressing six categories of emotions (anger, fear, disgust, happy, sad, and neutral). In Exp. 2, the morphing experiment, male voices expressing anger and fear were chosen to generate a set of morphed stimuli since these two voices have similar lengths. Anger to fear continua were created in eleven steps that corresponded to the fear/anger ratio as 0/100%, 10/90%, 20/80%, 30/70%, 40/60%, 50/50%, 60/40%, 70/30%, 80/20 %, 90/10%, and 100/0%. We used STRAIGHT for morphed stimuli generation (Kawahara et al., 2003). More specifically, voices were decomposed into five basic parameters by STRAIGHT: fundamental frequency (f_0_), frequency, duration, spectral-temporal density, and aperiodicity. Each parameter was manipulated independently. For each voice, we defined one time landmark and three frequency landmarks (corresponding to the first three formants) at the onset of the phonation and the same number of landmarks at the offset of the phonation. Morphed stimuli were generated based on the interpolation of these time-frequency landmark templates.

Stimuli were presented by MATLAB (The Mathworks, Natick, USA) using Psychophysics Toolbox 3.0 extension (Brainard, 1997), and were delivered via insert air-conduction earphones (ER2, Etymotic Research). Each stimulus was delivered 20 times and all stimuli were presented in a randomized order. For Exp. 1, a total of 240 trials (12 voices x 20 times) were presented. For Exp. 2, a total of 220 trials (11 morphed steps x 20 times) were presented. All stimuli were normalized to the same amplitude level. The sound volume was adjusted to the most comfortable level for each patient, approximately 65 dB sound pressure level (SPL). Patients were asked to press a button on a keyboard indicating which category of emotion she/he just heard using the hand ipsilateral to the side of the electrode’s coverage. No feedbacks were given to the subjects after each trial.

#### iEEG data analyses

We performed all data analyses using MATLAB. Each channel time series was visually inspected for artifacts. Channels with epileptiform activity were excluded from further analysis. Notch filters from Fieldtrip toolbox (http://www.fieldtriptoolbox.org/) were used to remove 50Hz noise and its second and third harmonics, and the data were down-sampled at 500Hz.

The data were then segmented with a 200-ms pre-stimulus baseline and a 1000-ms post-stimulus interval. We focused on the analysis of the high-gamma responses (70-140Hz), which have been shown to be highly correlated with functional magnetic resonance imaging (fMRI) blood-oxygen-level-dependent (BOLD) signals and population spike activities (Mukamel et al., 2005; Nir et al., 2007). The high-frequency broadband (HFB) responses of the high-gamma band were estimated using the following steps: (1) a 100-ms moving window (20-ms step) was used to perform short-time Fourier transform (STFT) for the pre-processed iEEG signal; (2) each frequency component time series was then normalized to its own baseline mean and was divided by its own baseline mean to determine its own HFB time series; (3) finally, HFB time series of each frequency component inside the range of 70-140 Hz were averaged together, producing a single time series for each trial of each channel. These procedures can cancel the 1/frequency decay of power in the spectrum.

In Exp. 1, the Emotion Recognition Experiment, patients that made wrong categorizations of the emotions in the emotion recognition task (the perceived emotion by the listener is not the same as the presented emotion by the speaker) were included in the analysis (see Supplementary Material for the details of subjects’ emotion recognition behavior results). The data analyses followed the pipeline showed in Figure S1. For each subject, electrode contacts located in HG, PI, AI, and amygdala were selected. HFB time series of all trials were calculated for each selected electrode. Presented emotion refers to the intended emotions expressed by the speaker, in which different types of emotion are determined by the acoustic properties of the voices. Perceived emotion refers to subject’s perceptual judgement of the voices as certain emotion types reflected in button pressing. To illustrate whether the responses of the electrodes were driven by the acoustic properties of the stimuli expressed by the speaker (presented emotion) or by the perceptual judgments of the listener (perceived emotion), we divided all trials into three stimulus-response patterns: “Match”, “Mismatch (False Alarm)”, and “Mismatch (Miss)”. “Match” referred to correct trials in which subject’s perceived emotion is the same as the presented emotion, i.e., emotion- *i* is being perceived as emotion- *i* (*i*|*i*); “Mismatch” referred to incorrect trials in which subject’s perceived emotion is different from the presented emotion. “Mismatch” trials were further divided into two error types relative to the “Match” trials, i.e., a “Mismatch (False Alarm)” trial had the same perceived emotion as a “Match” trial but with a different presented emotion (subject perceived emotion- *i* when other emotions were presented (*i*|*≠i*)), and a “Mismatch (Miss)” trial had the same presented emotion as a “Match” trial but with a different perceived emotion (emotion- *i* was presented but the subject perceived it as other emotions). To ensure that the “Mismatch” trials were not due to random mistakes made by subjects, subjects were expected to meet the following trial reassembling criteria: for a given presented emotion *i*, correctly categorize emotion *i* (“Match”) for more than 10 trials, categorize emotion *i* as other emotions (“Mismatch (Miss)”) for more than 10 trials, and categorize other emotions as emotion *i* (“Mismatch (False Alarm)”) for more than 10 trials. 16 out of the 24 subjects met all the criteria and were included for further data analysis (see Supplementary Material for the details of the three types of stimulus-response patterns from all subjects). In addition, only electrodes with significant responses (compared with baseline, p = 0.05 as the criterion, responsive electrodes) in at least one type of stimulus-response patterns (“Match”, “Mismatch (False Alarm)”, or “Mismatch (Miss)”) were included in further analyses, which resulted in 26 electrodes in HG, 11 electrodes in PI, 10 electrodes in AI, and 8 electrodes in amygdala. Since different stimuli have different durations which may induce different response onset and offset time, we only compared the maximum HFB responses across time of these three stimulus-response patterns in every single electrode. We conducted point-to-point two-sample t-test across trials on the time series of the HFB responses to the three types of trials, to determine the time window with significant differences (p < 0.05), and Bonferroni corrections were done across types. We defined a significant time window with at least 40-ms, and only time periods that have significant responses in at least one type of trials compared with baseline (two sample t-test, p < 0.05) were included in the statistical comparisons. We then defined stimulus selectivity index (SI_s_) and perception selectivity index (SI_p_) for a given “Match” condition ((*i*|*i*), emotion *i* was recognized as emotion *i*) as:

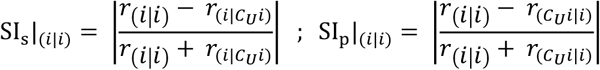

Where

U: set of all emotional stimuli; *r*_(*i*|*i*)_: maximim HFB of “Match” condition (*i*|*i*);

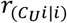: maximum HFB of “Mismatch(Miss)” condition in which stimulus is

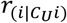: maximum HFB of “Mismatch(False Alarm)” condition in which response is *i*;

SI_s_ measures the normalized response differences between “Match” and “Mismatch (False Alarm)”, which reflects the selectivity to the acoustic properties of the stimuli (“Match” and “Mismatch (False Alarm)” trials have different presented emotions but same perceived emotions). SI_p_ measures the normalized response differences between “Match” and “Mismatch (Miss)”, which reflects the selectivity to the perceptual judgments of the perceivers (“Match” and “Mismatch (Miss)” trials have different perceived emotions but same presented emotions). Then 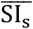 and 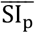 were derived by averaging SI_s_ and SI_p_ across all emotions respectively (only for emotions meet the trial reassembling criteria). Furthermore, we defined 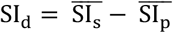 for single electrodes. 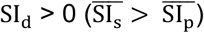 indicates the electrode is selective to the acoustic properties of the stimuli, while 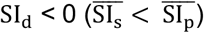 indicates the electrode is selective to subjective perceptual judgments of the perceivers. We conducted two-sample t-test to compare the SI_d_ between different brain areas (HG, PI, AI, and amygdala) across electrodes with Bonferroni corrections.

Latency of each brain area (HG, PI, AI, and amygdala) was calculated by averaging the peak latencies of HFB responses across all types of trials and all responsive electrodes in that brain area recorded from all patients.

To explore the discriminative latencies of AI and amygdala in discriminating different perceptual judgments, we calculated the d-prime values of responses to “Match” and “Mismatch (Miss)” stimuli at each time point in AI and amygdala for a single electrode as:

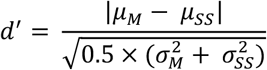

Where:

*μ*_*M*_: mean HFB of “Match” stimuli across trials

μ_*SS*_: mean HFB of “Mismatch(Miss)” stimuli across trials

σ_*M*_: standard deviation of HFB of “Match” stimuli across trials

σ_*SS*_: standard deviation of HFB of “Mismatch(Miss)” stimuli across trials

The d-prime value indicates the differences between responses to “Match” and “Mismatch (Miss)” stimuli. Then we conducted point-to-point paired t-test of the d-prime value at each point with the mean d-prime values at baseline across electrodes to find the significant time window.

To validate the robustness of the selectivity of single electrodes, we performed permutation tests by shuffling the trials of “Mismatch (False Alarm)” and “Mismatch (Miss)”. After running 1000 times of the permutation tests, a null distribution of SI_d_ was calculated. Significant level was determined by comparing the real value of SI_d_ with the null distribution.

In Morphing Experiment, to calculate the neural responses to morphed stimuli, we first calculated the psychometric function according to the emotion recognition behavior results of all subjects, and then HFB responses of all electrodes in response to the 11 steps of morphed stimuli were determined. 7 patients participated in this experiment and electrodes from the 7 patients that had significant (p<0.05, two-sample t-test) response differences to the original non-morphed anger (the fear/anger ratio as 0/100%) and fear (the fear/anger ratio as 100/0%) emotions were chosen for further analysis (5 electrodes in HG, 5 electrodes in PI, 4 electrodes in AI, 4 electrodes in amygdala). The HFB responses were normalized to the maximum response across all morphed stimuli so that we can compare the neural responses with behavior results directly. SSE (Sum of Squared Error) was calculated as:

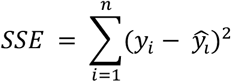

Where:

*n* : number of stimuli

*y*_*i*_ : Normalized HFB response to stimulus *i*

ŷ_*l*_ : predicted response to stimulus *i* (acoustic model or behavior result) SSE measures the distance between neural responses and predicted model.

## Results

### Responses of single electrodes in HG, PI, AI, and amygdala to emotional voices

Figure 1A, B shows the locations of all electrodes in HG, PI, AI, and amygdala implanted in all subjects by aligning them in an average brain. The parcellation of PI and AI were defined using the resting state functional magnetic resonance imaging (rsfMRI) data in each subject as previously described (Zhang et al., 2019). Figure 1D shows high-frequency broadband (HFB, See Methods) responses of iEEG signals to an emotional stimulus (“Disgust 2”) recorded from electrodes in HG, PI, AI, and amygdala, respectively (Figure 1C). Responses in HG and PI showed higher response amplitudes and shorter peak latencies than responses in AI and amygdala (Figure 1D), consistent with our previous findings that PI and AI have distinct functions (Zhang et al., 2019).

### Reassemble response trials based on the presented emotions of stimuli and the perceived emotions of listeners

To explore whether insular responses to emotional stimuli were correlated with acoustic properties of the stimuli (the presented emotions) or a listener’s subjective perception of the emotional content of the stimuli (the perceived emotions), we regrouped the response trials according to the emotion recognition behavior task results. “Match” trials were correct trials where the perceived emotion is the same as the presented emotion of the stimulus, i.e., emotion- *i* is being perceived as emotion- *i* (*i*|*i*). Incorrect trials were divided into two types: “Mismatch (False Alarm)” trials - incorrect trials where the subject perceived emotion- *i* when other emotions were presented (*i*|≠ *i*). “Mismatch (Miss)” trials-incorrect trials where emotion- *i* was presented but the subject perceived it as other emotions (≠ *i*|*i*).

For example, when a subject was presented with a sound with emotion “anger”, if the subject perceived the emotion as “anger”, this trial was considered a “Match” trial (Figure 2A, brown color). However, if the subject perceived it as other emotions, this trial was considered a “Mismatch (Miss)” trial (Figure 2A, green color). On the other hand, if the subject perceived the emotion “anger” when presented with other emotions, the trial was considered as a “Mismatch (False Alarm)” trial (Figure 2A, yellow color). Table 2 shows the trial regroup results from all subjects. 16 out of the 24 subjects met the trial reassembling criteria and were included for further data analysis. Please also see Supplementary Material for more details. Based on this analysis framework, if similar neural responses are observed between “Match” and “Mismatch (False Alarm)” trials in an electrode, it suggests that the responses of this electrode are primarily driven by the perceived emotion. On the other hand, if similar responses are observed between “Match” and “Mismatch (Miss)” trials in an electrode, it suggests that the responses in this electrode are primarily driven by presented emotion of the stimuli. iEEG data were analyzed by averaging HFB responses from each electrode according to the three types of trials (at least 10 trials in each type). Figure 3 shows representative HFB responses from four representative electrodes in HG, PI, AI, and amygdala, respectively, analyzed across the three types of trials. The electrode in HG (Figure 3A) showed similar response profiles among “Match”, “Mismatch (False Alarm)”, and “Mismatch (Miss)” conditions between 0 and 400ms after the stimulus onset, with peak latencies of the responses at around 170ms. No significant differences of peak responses were observed among the three types of trials in this electrode (Figure 3A, inset, p > 0.05, two-sample t-test with Bonferroni correction across types), suggesting that the responses of this HG electrode were not dependent on the presented or perceived emotions. The electrode in PI (Figure 3B) responded similarly in “Match” and “Mismatch (Miss)” conditions (0-300ms after stimulus onset) with peak responses at around 170ms which is similar to that of HG electrode (Figure 3A), but with a much lower response in “Mismatch (False Alarm)” condition. There was no significant difference between “Match” and “Mismatch (Miss)” conditions but a significant difference between “Match” and “Mismatch (False Alarm)” conditions (Figure 3B, grey bar and inset, p = 0.05 is the criterion, two-sample t-test with Bonferroni correction across types), suggesting that responses of this PI electrode were largely driven by presented emotions of the stimuli, independent of a listener’s subjective perception of the emotions embedded in the voices. In contrast, the electrode in AI (Figure 3C) responded similarly between “Match” and “Mismatch (False Alarm)” conditions that were significantly different from “Mismatch (Miss)” condition (Figure 3C, grey bar and inset, p = 0.05 is the criterion, two-sample t-test with Bonferroni correction across types), suggesting that responses of this AI electrode were primarily driven by the perceived emotion. Note that the responses peaked at around 220ms which is substantially later than those of HG and PI electrodes (Figure 3A, B). The electrode in amygdala (Figure 3D) also responded similarly between “Match” and “Mismatch (False Alarm)” conditions with peak responses at around 320ms, but significantly different from “Mismatch (Miss)” condition (Figure 3D, grey bar and inset, p = 0.05 is the criterion, two-sample t-test with Bonferroni correction across types). The examples in Figures 3C and 3D show that the responses of AI and amygdala electrodes are largely driven by a listener’s subjective perception of the emotions embedded in the voices.

**Figure 2.**
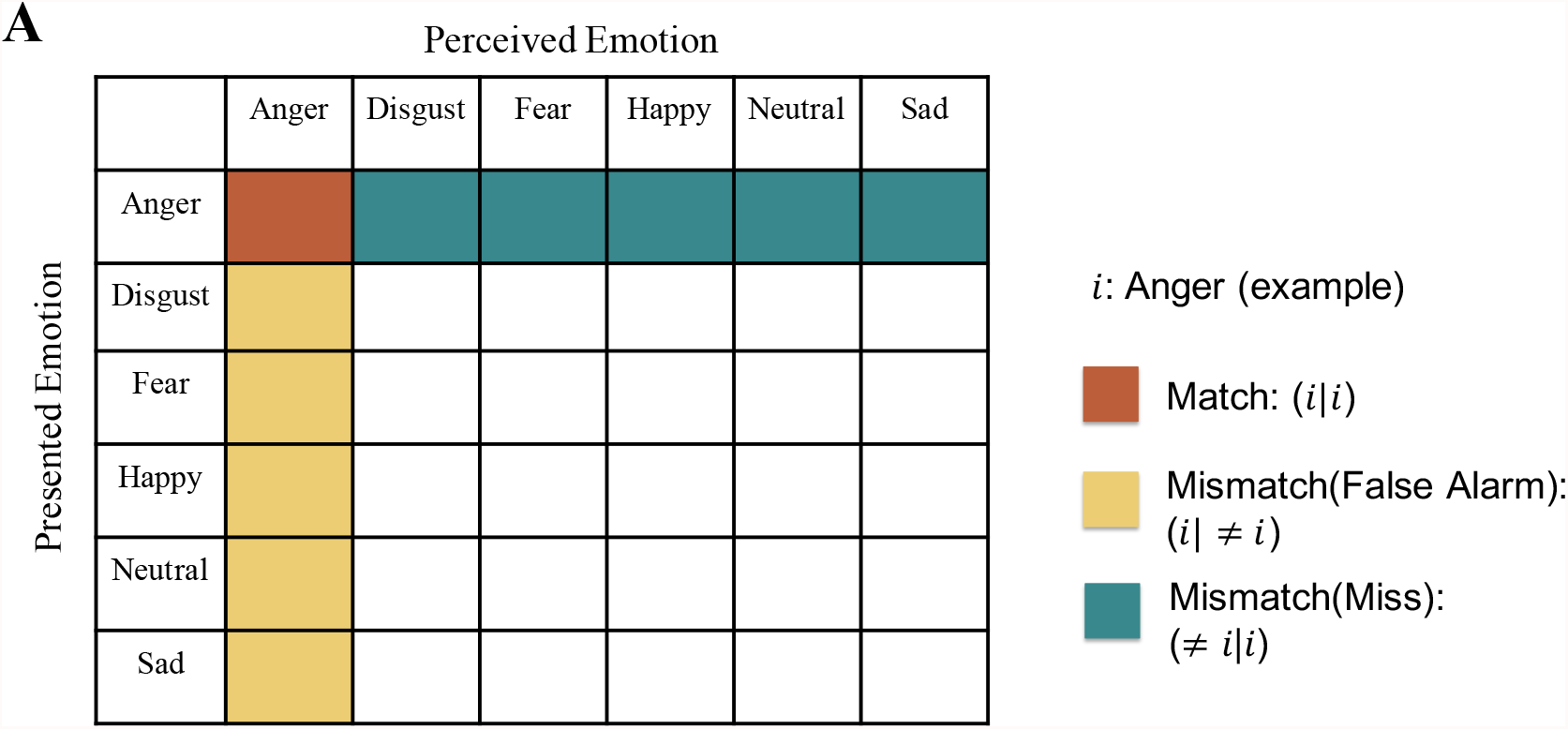
Illustration of trial reassembling method for presented “Anger”: (A) Example of trial reassembling method. Trials are divided into three types: Match, Mismatch (False Alarm), and Mismatch (Miss) according to the consensus emotions of the stimuli (presented emotion) and emotion recognition behavior results (perceived emotion). Match: emotion *i* recognized as emotion *i* (*i*|*i*) (brown rectangle). Mismatch (False Alarm): emotions other than *i* recognized as emotion *i* (*i*| ≠ *i*) (yellow rectangle). Mismatch (Miss): emotion *i* recognized as emotions other than *i* (≠ *i*|*i*) (green rectangle). Left table shows the three categories under an example of emotion *i* (emotion *i* = anger). For each emotion type, two voices (one from male and the other from female) with 20 repetitions were included, resulting in 40 trials. Only conditions that have Match trials exceeded 10, Mismatch (False Alarm) trials exceeded 10, and Mismatch (Miss) trials exceeded 10 were included for further analysis.

**Table 2.** Trial regroup results of all subjects

| Subject | Condition number that matches the trial regroup criterion | Presented emotion type for all matched conditions | “Match” trial number | “Mismatch (Miss)” trial number | “Mismatch (False Alarm)” trial number |
| --- | --- | --- | --- | --- | --- |
| S1 | 2 | Anger<br>Neutral | 12<br>25 | 28<br>15 | 41<br>14 |
| S2 | 0 |  |  |  |  |
| S3 | 1 | Disgust | 11 | 29 | 16 |
| S4 | 1 | Disgust | 17 | 23 | 33 |
| S5 | 0 |  |  |  |  |
| S6 | 1 | Neutral | 12 | 28 | 34 |
| S7 | 0 |  |  |  |  |
| S8 | 2 | Anger<br>Disgust | 10<br>20 | 30<br>20 | 41<br>55 |
| S9 | 1 | Disgust | 18 | 22 | 44 |
| S10 | 1 | Anger | 18 | 22 | 12 |
| S11 | 0 |  |  |  |  |
| S12 | 0 |  |  |  |  |
| S13 | 1 | Fear | 29 | 11 | 22 |
| S14 | 3 | Anger | 14 | 26 | 17 |
|  |  | Disgust | 11 | 29 | 31 |
|  |  | Fear | 19 | 21 | 18 |
| S15 | 0 |  |  |  |  |
| S16 | 1 | Sad | 20 | 20 | 38 |
| S17 | 1 | Sad | 20 | 20 | 119 |
| S18 | 0 |  |  |  |  |
| S19 | 2 | Disgust | 27 | 13 | 34 |
|  |  | Fear | 17 | 23 | 27 |
| S20 | 3 | Anger | 16 | 24 | 39 |
|  |  | Fear | 25 | 15 | 11 |
|  |  | Neutral | 10 | 30 | 17 |
| S21 | 1 | Fear | 13 | 27 | 61 |
| S22 | 0 |  |  |  |  |
| S23 | 2 | Disgust | 26 | 14 | 41 |
|  |  | Fear | 24 | 16 | 36 |
| S24 | 1 | Fear | 15 | 25 | 43 |

**Figure 3.**
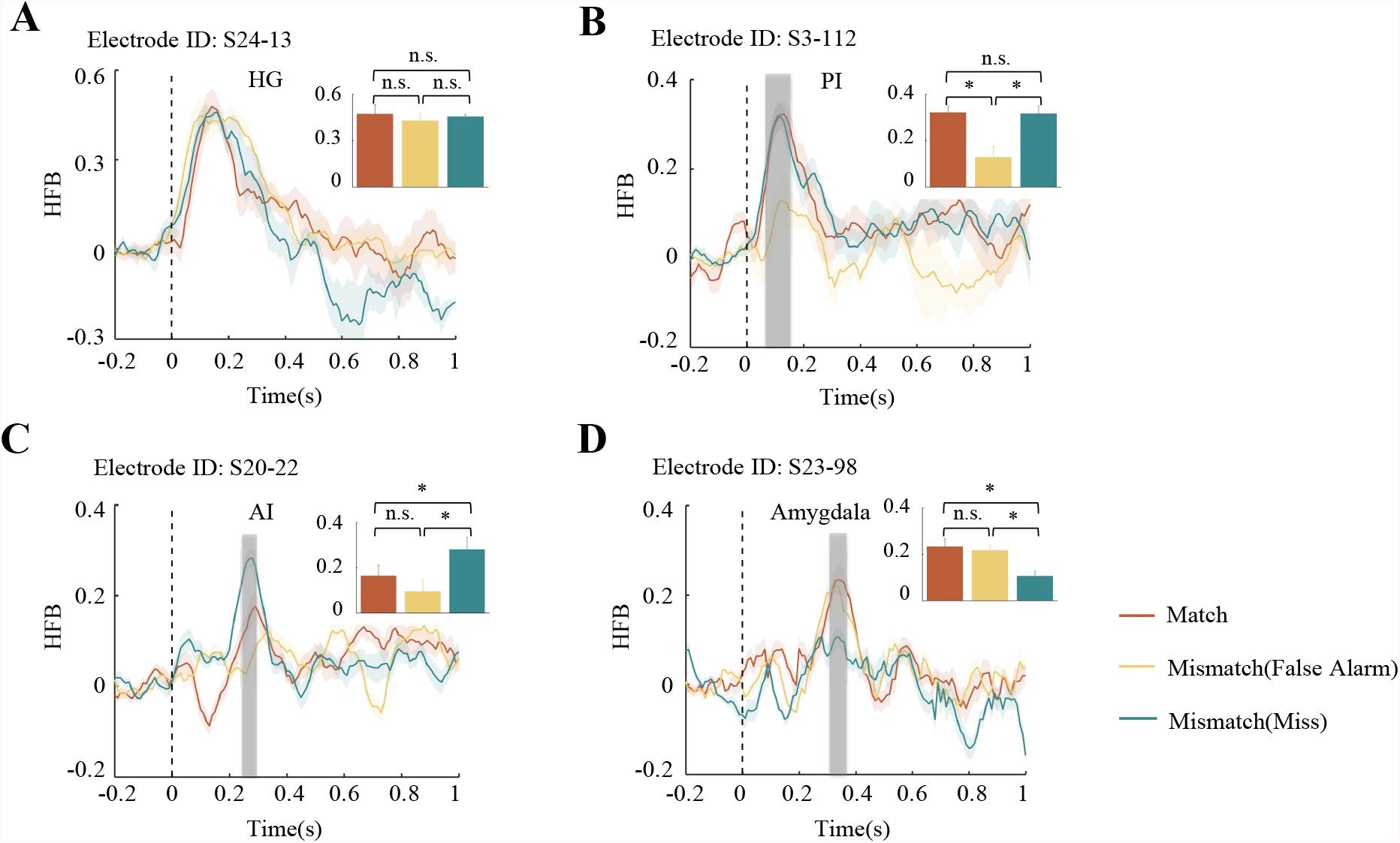
Average HFB time series of single electrode examples across the three types of trials (Match, Mismatch (False Alarm), and Mismatch (Miss)): (A) Example electrode in HG. (B) Example electrode in PI. (C) Example electrode in AI. (D) Example electrode in amygdala. (mean ± sem; grey region: statistically significant time window across types around the response peaks, only computed for time window with significant responses to at least one type of trials compared with baseline; vertical black dash line: stimulus onset) Inset panels: Comparisons of HFB responses at peak points across trials. (n.s.: not significant; *: p < 0.05; two-sample t-test with Bonferroni correction)

### Population properties of the neural activities that biased toward the presented emotions or the perceived emotions

To quantify population properties across all electrodes (26 electrodes in HG, 11 electrodes in PI, 10 electrodes in AI, 8 electrodes in amygdala), we calculated stimulus selectivity index (SI_s_) and perception selectivity index (SI_p_) for each emotional stimulus in each electrode. SI_s_ measures the normalized response difference between “Match” and “Mismatch (False Alarm)” conditions for a given emotional stimulus, whereas SI_p_ measures the normalized response difference between “Match” and “Mismatch (Miss)” conditions for a given emotional stimulus. 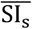 and 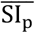 were computed by averaging SI_s_ and SI_p_ across all conditions (for emotions meet the trial reassembling criteria) in each electrode, respectively. Figure 4A plots 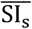 versus 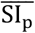 for all electrodes in HG, PI, AI, and amygdala. The inset panel in Figure 4A shows the percentages of electrodes with 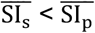 or 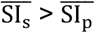 in each brain region. We further defined 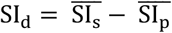 to quantify whether the response of an electrode is associated with acoustic properties of the stimuli (the presented emotions) or is associated with subjective perception of the embedded emotion (the perceived emotions). SI_d_> 0 indicates that the electrode’s response is biased towards acoustic properties of the stimuli, whereas SI_d_< 0 indicates that the electrode’s response is biased towards subjective perception of the embedded emotion. Figure 4B shows SI_d_ values averaged across all electrodes in HG, PI, AI, and amygdala, respectively. Note that HG and PI have positive SI_d_ values, indicating a bias towards acoustic properties of the stimuli, while AI and amygdala have negative SI_d_ values, indicating a bias towards subjective perception of the embedded emotion. The difference between SI_d_ values of HG and PI and those of AI and amygdala are significant (p < 0.01; two-sample t-test with Bonferroni correction). To validate the robustness of the response properties of single electrodes, we conducted permutation tests by shuffling the trials of “Mismatch (False Alarm)” and “Mismatch (Miss)”. SI_d_ null distributions were derived by permuting 1000 times. Figure 5 shows the comparisons between original SI_d_ values and the null distributions. SI_d_ values of HG, PI, AI, and amygdala are all significantly different from null distributions (p < 0.05). All together, these results indicate that responses of HG and PI are associated with acoustic properties that determined the presented emotion of the stimuli, whereas responses of AI and amygdala are largely biased toward the subjective perception of emotions.

**Figure 4.**
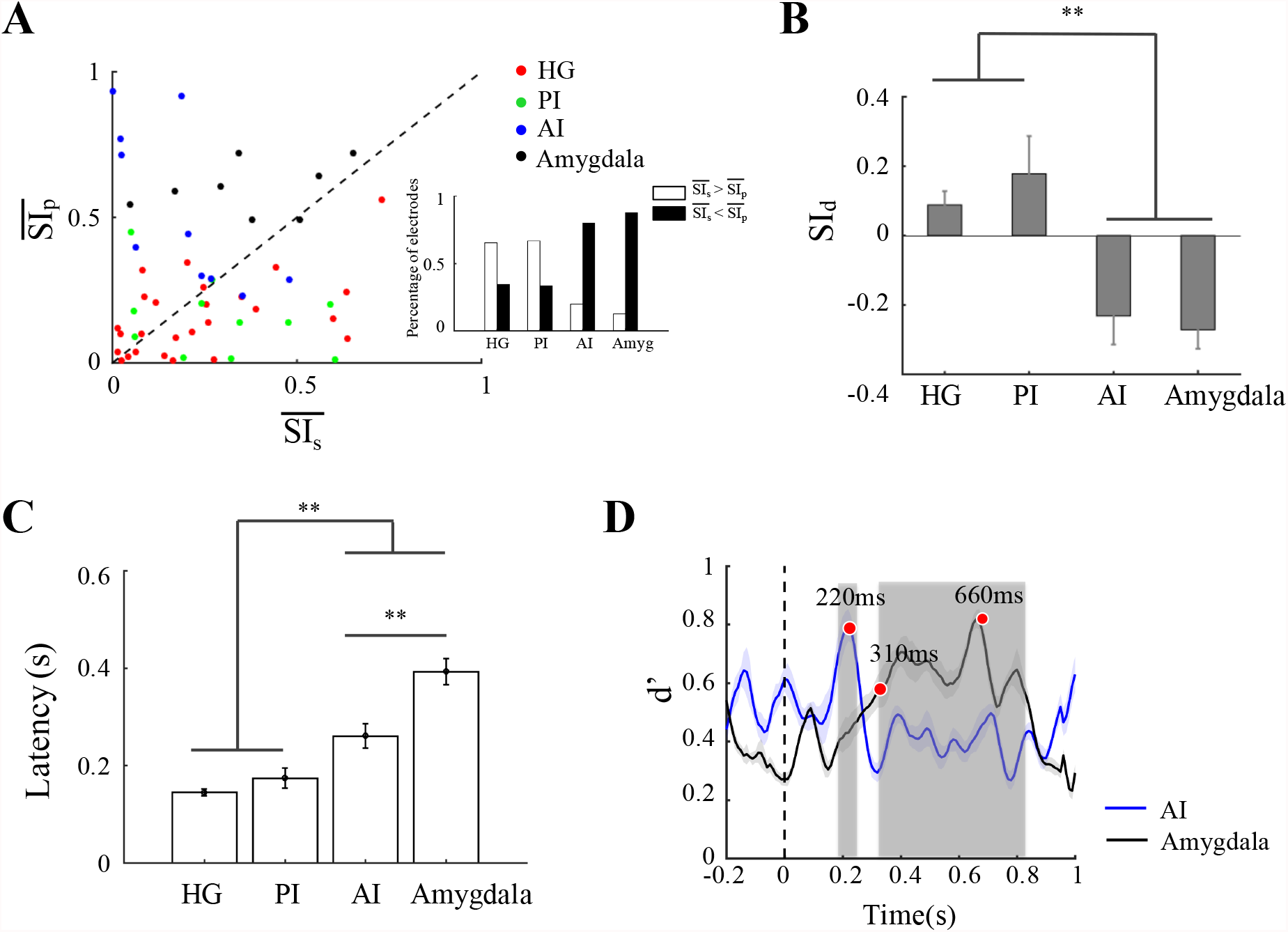
Neural activities following objective acoustic properties or subjective perceptions: (A) 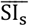 and 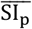 distribution across all electrodes in HG, PI, AI, and amygdala. Inset panel: percentage of electrodes with 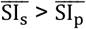 and 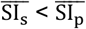. (B) SI_*d*_ in HG, PI, AI, and amygdala across all electrodes (mean ± sem; **: p < 0.01; two-sample t-test with Bonferroni correction). (C) Peak latencies of HFB responses averaged across all types of stimuli and electrodes in each region (mean ± sem; **: p < 0.01; two-sample t-test with Bonferroni correction). (D) Discriminative latencies, d-prime values (mean ± sem across electrodes; shadow area: statistically significant area; compared with baseline) of responses to Match and Mismatch (Miss) stimuli across time in AI and amygdala.

**Figure 5.**
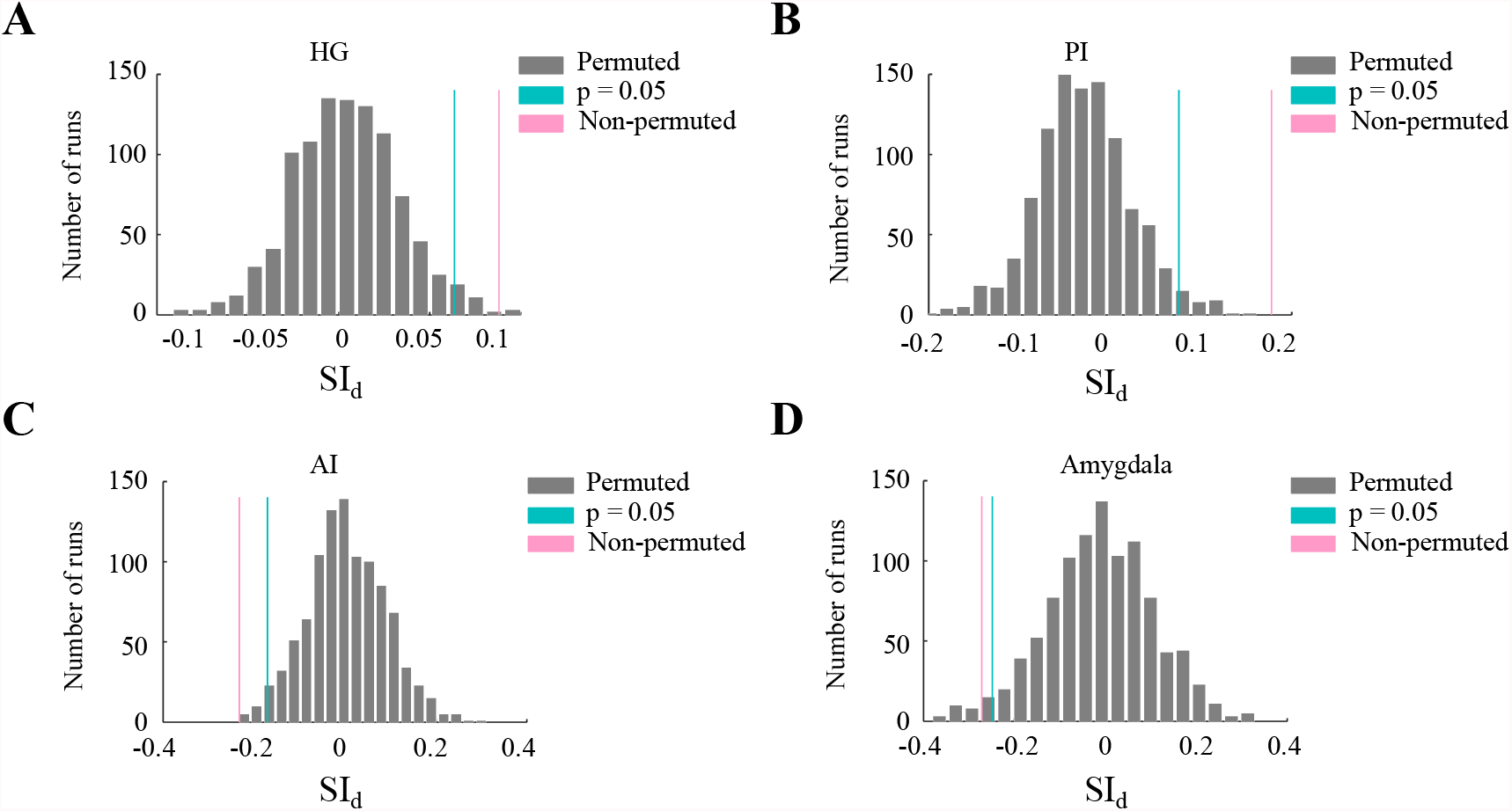
Control analyses, using population summary metrics, compared with permutation tests: (A) All electrodes in HG. (B) All electrodes in PI. (C) All electrodes in AI. (D) All electrodes in amygdala. The gray distributions are the null distributions from permutation tests. The green bars are the significant levels, and the red bars are the population summary metrics across all electrodes in that brain region.

Figure 4C compares peak latencies between the four brain regions, averaged over all stimuli and electrodes and shows a trend of successively increasing latency from HG to PI, AI, and amygdala. Latencies of HG and PI are significantly different from those of AI and amygdala, and amygdala has a significantly longer latency than AI (Figure 4C). To further understand the time courses of the selectivity to subjective perceptions in AI and amygdala, we calculated the d’ values of each time point across electrodes in AI and amygdala, to compare the latencies of AI and amygdala in discriminating different subjective perceptions (discrimination of “Match” and “Mismatch (Miss)”) (Figure 4D). Grey regions show the time window with significant differences between the d’ value at the time point and mean d’ value at baseline across electrodes. We found that AI responses significantly discriminated different subjective perceptions at around 220ms after stimulus onset, while amygdala showed significant discrimination between 310-820ms (Figure 4D). The latency data suggest that the representation of acoustic stimuli is transformed to the representation of perceived emotions through a hierarchical pathway, in which insula may function as a bridge between HG and amygdala.

### Neural response properties to morphed stimuli

In light of the above results, we conducted a Morphing Experiment. We tested in some subjects (7 subjects participated in the morphing experiments) how HG, PI, AI, and amygdala responded to a continuum of stimuli morphed between two emotional stimuli (Table 1). A “fear” to “anger” continuum was generated by linearly morphing the two stimuli in eleven equal steps (see Methods) (Figure 6A). The subjects were asked to identify if a stimulus was perceived as “fear” or “anger”. Their performance, averaged across all subjects, showed a sigmoid-like psychometric curve (Figure 6B), indicating a categorical pattern to these stimuli. iEEG responses were recorded from HG, PI, AI, and amygdala at each step of the morphed stimuli while the subjects performed the tasks. Figures 6C-F show representative HFB responses from the four brain regions and comparisons between these responses and acoustic (Figure 6A) or behavior (Figure 6B) curves across the stimulus continuum. Electrodes with significant differences between the responses to non-morphed “anger” and “fear” stimuli were selected for this analysis (5 electrodes in HG, 5 electrodes in PI, 4 electrodes in AI, 4 electrodes in amygdala). The left plots of Figures 6C-F show HFB time series for each stimulus of the continuum. HFB response magnitude for each stimulus was calculated by averaging over the time window indicated by the grey rectangles on Figure 6C-F left plots (significant HFB response time window compared with baseline responses with a duration less than 200ms) and normalized to values between 0 and 1. The right plots of Figure 6C-F show the comparisons between normalized HFB response magnitudes and acoustic (red solid line) and behavior (red dash line) curves across the stimulus continuum for each representative electrode. We calculated SSE (Sum of Squared Error, see Methods) to quantify the distance between the normalized HFB response magnitude and acoustic or behavior curve as a function of morph steps (Figure 6C-F, right plot inset). A higher SSE value indicates a larger distance between two measures. We found that the normalized HFB response magnitudes of the electrodes in HG and PI are closer to the acoustic curve than the behavior curve (Figure 6C, D right plot inset), suggesting that neural responses in these two regions are associated with acoustic properties that determined the presented emotions of the stimuli. In contrast, the normalized HFB response magnitudes of the electrodes in AI and amygdala are closer to the behavior curve than the acoustic curve (Figure 6E, F right plot inset), suggesting that neural responses in these two regions are biased toward the perceived emotions. These results agree with our observations from Exp. 1. To demonstrate the population characteristics of these observations, we calculated the SSE of all electrodes in each brain region. Figure 6G shows the average SSE values across all electrodes in HG, PI, AI, and amygdala, respectively. HG and PI have significantly smaller SSE values for acoustic than SSE values for behavior (p < 0.05; paired t-test with Bonferroni correction), whereas AI and amygdala have significantly smaller SSE values for behavior than SSE values for acoustic (p < 0.05; paired t-test with Bonferroni correction). These results support the observations that responses in HG and PI are mostly determined by acoustic properties of the stimuli, whereas the responses in AI and amygdala are largely determined by perceived emotions.

**Figure 6.**
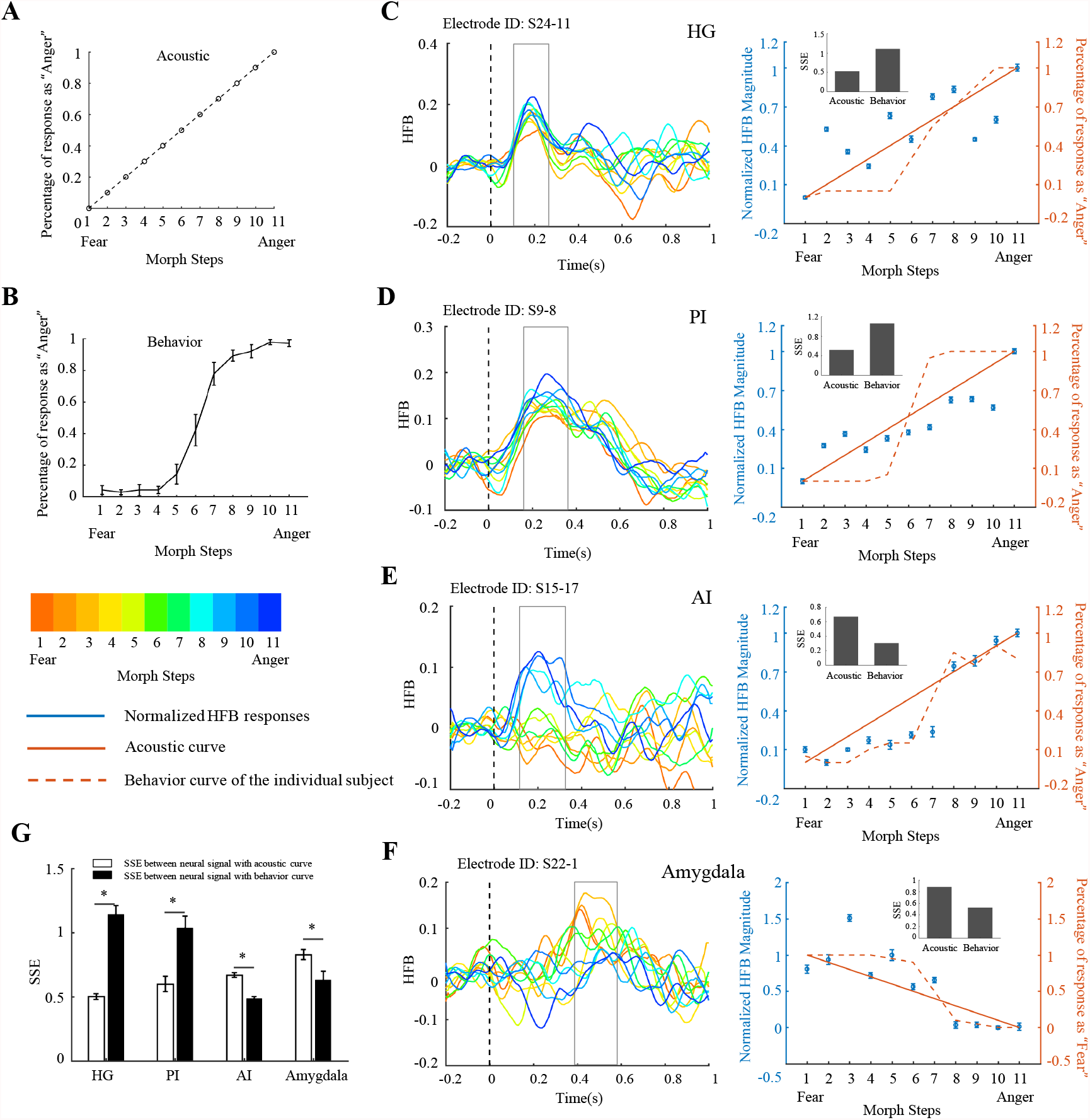
Activation patterns to morphed stimuli: (A) Theoretical psychometric curve based on acoustic features to all morphed stimuli (acoustic psychometric curve). (B) Real psychometric curve to all morphed stimuli averaged across all subjects (behavior psychometric curve). (C - F) Left panel: HFB responses to all morphed stimuli across time of representative electrodes in HG (C), PI (D), AI (E), and amygdala (F); Right panel: Comparisons among normalized HFB response magnitudes (responses were averaged across the time window indicated by grey rectangle region in left panel and normalized to the maximum responses, blue color) of the example electrodes, acoustic psychometric curve (red solid line), and behavior psychometric curve of the subject implanted with the example electrode (red dash line), inset plot: SSE (sum of squares errors) between normalized HFB magnitude and acoustic curve as well as between normalized HFB magnitudes and behavior curve. (G) Comparisons of SSE between normalized HFB response magnitudes with acoustic psychometric curve and with behavior psychometric curve across all electrodes in HG, PI, AI, and amygdala (*: p < 0.05; paired t-test with Bonferroni correction).

## Discussion

### Auditory emotion recognition pathways

In humans, auditory emotion recognition is thought to be accomplished through a hierarchical model (Schirmer and Kotz, 2006; Morningstar et al., 2018). This model consists of two steps. First, low-level processing of acoustic properties for categorizing auditory objects and extracting emotional contents occurs in HG (Wildgruber et al., 2006), the “temporal voice areas” (TVA) in lateral superior temporal gyrus (STG, Yovel and Belin, 2013), and bilateral STG. Second, higher-level cognition and evaluation of emotional contents occurs in anterior cingulate cortex (ACC), amygdala, insula (Bestelmeyer et al., 2014), and dorsomedial and lateral frontal cortex (Schirmer and Kotz, 2006). However, most of the neuroimaging studies in support of this model used subtraction designs, which make it difficult to demonstrate the existence of separate steps and to reveal the different functions of HG, STG, amygdala, insula, ACC, and frontal cortex during auditory emotion recognition tasks. In this study, we recorded iEEG signals from epilepsy patients implanted with depth electrodes in the insula (PI and AI regions), amygdala, and HG. iEEG signals have much higher temporal resolution than neuroimaging signals. Thus, we are able to compare the response properties and response latencies of these four regions during the auditory emotion recognition tasks. We focused on the analyses of electrodes from HG, insula, and amygdala due to the limited electrode coverage of all the patients included in this study. The roles of STG, ACC, and frontal cortex in auditory emotion recognition are awaited for further investigations.

Increased functional connectivity between HG and insula during emotion recognition tasks (Karle et al., 2018) as well as decreased functional connectivity between HG and insula in psychiatric conditions (schizophrenia) with perceptual deficits for recognizing emotions embedded in human voice (Kantrowitz et al., 2015) were observed, suggesting critical roles of human insula for auditory emotion recognition. In addition, previous studies have shown that posterior insula is structurally and functionally connected to auditory cortex, while anterior insula is structurally and functionally connected to the amygdala (Cauda et al., 2011; Zhang et al., 2019). The present study provided new insights in different roles that posterior and anterior insula may play in processing emotional information embedded in human voices. We showed that neural responses in anterior insula and amygdala depend on subjective perception of the emotional contents embedded in voices, whereas neural responses in posterior insula and Heschl’s gyrus are associated with the acoustic properties of voices. In addition, we observed a progressive increase in response latency from HG to posterior insula, anterior insula, and to amygdala. Based on these observations, we propose that human insula functions as a bridge to link objective acoustic information processed in auditory cortex to subjective emotion perception in amygdala. Previous studies have proposed a dual-brain-pathways model, which consists of a ventral pathway and a dorsal pathway, for processing speech, both starting from auditory cortex. In this model, the ventral pathway consists of structures in the anterior and middle portions of the temporal lobe and is involved in speech comprehension; the dorsal pathway consists of structures in the frontal lobe, the posterior dorsal-most portions of the temporal lobe, and the parietal operculum, and is involved in translating acoustic speech signals into articulatory representations (Rauschecker and Scott, 2009; Hickok and Poeppel, 2007). Our findings suggest a potential auditory emotion recognition pathway leading from auditory cortex to the posterior insula, anterior insula, and then amygdala for transforming external acoustic information to internal perceived emotion information.

### Multimodal convergent functional architecture of human insula

Previous studies (Cauda et al., 2011; Zhang et al., 2019) of structural and functional connectivity using diffusion MRI and rsfMRI showed that human insula is involved in two distinct neural networks: 1) the posterior insula is connected to sensorimotor, premotor, supplementary motor, olfactory, temporal cortex, and some occipital areas; 2) the anterior insula is connected to the anterior cingulate cortex, the middle and inferior frontal cortex, the amygdala, and the temporoparietal cortex. The connectivity patterns of insula provide neural basis for its multimodal convergent functions.

Functional imaging and electrical stimulation studies in human and non-human primate have suggested that the insula is involved in processing sensorimotor (Ostrowsky et al., 2002), auditory (Bamiou et al., 2003; Remedios et al., 2009; Zhang et al., 2019), visual (Augustine, 1996), pain (Brooks et al., 2005; Baliki et al., 2009; Schweinhardt et al., 2006; Wager et al., 2013), gustatory, olfactory (Shelley and Trimble, 2004), affective (Barrett and Wager, 2006; Caruana et al., 2011; Zaki et al., 2012), interoceptive (Craig, 2002, 2009), and high-level cognitive information (Sridharan et al., 2008). These suggest that insula functions as an important site of multimodal convergence, involving in sensory processing as well as higher cognitive functions. Therefore, the transformation of external sensory representation to internal subjective emotional representation from PI to AI found in this study may also apply to other sensory modalities, that sensory information with emotional contents from all sensory modalities can be first conveyed from sensory cortex to the PI, and emotional information is then extracted and perceived in the AI and sent to the amygdala and frontal cortex. However, further studies are needed to elucidate the underlying neural mechanisms that how the transformation from PI to AI take place.

### Functional role of AI in emotion perception

Recognizing the emotional states of others and experiencing the emotional states by oneself are two different emotion perception processes. However, recent studies have suggested shared emotional network of the two processes, that regions of the human brain are activated similarly to an emotional state affecting oneself or others. For example, Wicker et al. showed that AI is engaged not only in experiencing disgusting/pleasing tastes and odors, but also in viewing disgusted/pleased faces (Wicker et al., 2003). In addition, direct pain experience activates a network including insula, cingulate, and somatosensory areas. Among those regions, especially insula and cingulate cortex, are also activated when observing pain delivered to others (Singer et al., 2004; Lamm et al., 2011). There observations have been interpreted as emotional empathy and they all highlight the importance of AI in the two processes (Caruana et al., 2011). Roles of human insula in conscious perceptions and awareness have also been reported (Craig, 2002, 2009). Findings of the present study that AI is selective to the perceived emotions suggest that recognizing the emotional states of others requires inner representations but not depends on sensory stimulus itself. This is consistent with the concept that recognizing the emotional states by others is an inner process (Craig, 2002, 2009). To conclude, our results further confirm the role of AI in recognizing emotional states of others.

Another intriguing question is whether different emotions are being processed differently in AI and amygdala. Previous studies have demonstrated the non-uniformed representation of emotions in amygdala, with prominent responses to fear-related aversive stimuli (Adolphs et al., 1997; LeDoux, 2017; Davis and Whalen, 2001). However, some studies also provide the evidence showing the role of amygdala in positive emotion processing (Adolphs, 2010, Anderson and Adolphs, 2014; Hamann et al., 1999). In addition, studies have showed both positive and negative emotions can activate AI (Craig, 2002, 2009). These studies indicate both AI and amygdala can be driven by multiple emotions.

However, there is no evidence showing whether the representations of multiple emotions are different in AI and amygdala. In this study, we attempted to compare the responses to different emotions in AI and amygdala by using classification methods. However, no differences were observed. There are two possible explanations. First, Different emotions may be represented in AI and amygdala by different spatial patterns, however, the iEEG electrodes implanted in our patients have large sizes that lack spatial resolution to differentiate responses from different emotional contents. Second, different emotions may be represented by the different connectivity patterns among AI, amygdala, and other high-level cortical areas (such as frontal cortex, cingulate cortex), that comparing the responses in one area is not sufficient to uncover the differences. Future studies using higher spatial resolution recordings (such as high density iEEG electrodes) and covering more cortical areas may have the potential to solve this question.

## Author Contributions

Y.Z., J.H., B.H. and X.W. conceived the experiments. Y.Z. and J.H. selected the emotional stimuli and designed the experimental paradigm. Y.Z. performed experiments. W.J.Z. performed the neurosurgeries on the epilepsy patients. Y.Z. and X.W. analyzed and interpreted data. Y.Z., J.H. and X.W. wrote the manuscript.

## Acknowledgments

We thank Y. Ding, H. Han, X.P. Si, C. Song, D. Zhang and Y.L. Yan for comments and discussion. This work was supported by a grant from National Science Foundation of China (No. 61473169) to B.H..

## Data availability

The data that support the findings of this study are available from the corresponding author upon reasonable request.

## Code availability

Data analysis was performed with custom MATLAB code, available upon reasonable request from the corresponding author.

## References

Adolphs R (2010) What does the amygdala contribute to social cognition? Ann. N. Y. Acad. Sci. 1191: 42–61.

Adolphs R, Cahill L, Schul R, Babinsky R (1997) Impaired declarative memory for emotional material following bilateral amygdala damage in humans. Learning & Memory. 4: 291–300.

Anderson DJ, Adolphs R (2014) A framework for studying emotions across species. Cell. 157: 187–200.

Augustine JR (1985) The insular lobe in primates including humans. Neurol. Res. 7: 2–10.

Augustine JR (1996) Circuitry and functional aspects of the insular lobe in primates including humans. Brain Res. Rev. 22: 229–244.

Baliki MN, Geha PY, Apkarian AV (2009) Parsing pain perception between nociceptive representation and magnitude estimation. Journal of neurophysiology. 101: 875–887.

Bamiou DE, Musiek FE, Luxon LM (2003) The insula (Island of Reil) and its role in auditory processing: literature review. Brain Res. Rev. 42: 143–154.

Barrett LF, Wager TD (2006) The structure of emotion evidence from neuroimaging studies. Curr Dir Psychol Sci. 15:79–83.

Belin P, Fillion-Bilodeau S, Gosselin F (2008) The Montreal Affective Voices: a validated set of nonverbal affect bursts for research on auditory affective processing. Behav. Res. Methods. 40: 531–539.

Bestelmeyer PE, Maurage P, Rouger J, Latinus M, & Belin P (2014) Adaptation to vocal expressions reveals multistep perception of auditory emotion. Journal of Neuroscience. 34: 8098–8105.

Brainard DH (1997) The psychophysics toolbox. Spatial vision. 10: 433–436.

Brooks JC, Zambreanu L, Godinez A, Tracey I (2005) Somatotopic organisation of the human insula to painful heat studied with high resolution functional imaging. Neuroimage. 27: 201–209.

Brugge JF, Nourski KV, Oya H, Reale RA, Kawasaki H, Steinschneider M, Howard III MA (2009) Coding of repetitive transients by auditory cortex on Heschl’s gyrus. Journal of neurophysiology. 102: 2358–2374.

Caruana F, Jezzini A, Sbriscia-Fioretti B, Rizzolatti G, Gallese V (2011) Emotional and social behaviors elicited by electrical stimulation of the insula in the macaque monkey. Curr Biol. 21:195–199.

Cauda F, D’Agata F, Sacco K, Duca S, Geminiani G, Vercelli A (2011) Functional connectivity of the insula in the resting brain. Neuroimage. 55: 8–23.

Chang EF, Rieger JW, Johnson K, Berger MS, Barbaro NM, Knight RT (2010) Categorical speech representation in human superior temporal gyrus. Nature neuroscience. 13: 1428.

Craig AD (2002) How do you feel? Interoception: the sense of the physiological condition of the body. Nat. Rev. Neurosci. 3: 655–666.

Craig AD (2009) How do you feel--now? The anterior insula and human awareness. Nat. Rev. Neurosci. 10: 59–70.

Davis M, Whalen PJ (2001) The amygdala: vigilance and emotion. Molecular psychiatry. 6: 13–34.

Ding Y, Zhang Y, Zhou W, Ling Z, Huang J, Hong B, & Wang X (2019) Neural correlates of music listening and recall in the human brain. J. Neurosci. 39: 8112–8123.

Ellis, Andrew W. “Neuro-cognitive processing of faces and voices.” Handbook of research on face processing. (Elsevier, 1989), pp. 207–215.

Fitzgerald DA, Angstadt M, Jelsone LM, Nathan PJ, & Phan KL (2006) Beyond threat: amygdala reactivity across multiple expressions of facial affect. Neuroimage. 30: 1441–1448.

Griffiths TD, Kumar S, Sedley W, Nourski KV, Kawasaki H, Oya H, … & Howard MA (2010) Direct recordings of pitch responses from human auditory cortex. Current Biology. 20: 1128–1132.

Hamann SB, Ely TD, Grafton ST, Kilts CD (1999) Amygdala activity related to enhanced memory for pleasant and aversive stimuli. Nature neuroscience. 2: 289–293.

Hickok G, Poeppel D (2007) The cortical organization of speech processing. Nat. Rev. Neurosci. 8: 393–402.

Höistad M, Barbas H (2008) Sequence of information processing for emotions through pathways linking temporal and insular cortices with the amygdala. Neuroimage. 40: 1016–1033.

Kantrowitz JT, Hoptman MJ, Leitman DI, Moreno-Ortega M, Lehrfeld JM, Dias E, … & Javitt DC (2015) Neural substrates of auditory emotion recognition deficits in schizophrenia. Journal of Neuroscience. 35: 14909–14921.

Karle KN, Ethofer T, Jacob H, Brück C, Erb M, Lotze M, … & Kreifelts B (2018) Neurobiological correlates of emotional intelligence in voice and face perception networks. Social cognitive and affective neuroscience. 13: 233–244.

Kawahara, Hideki, Hisami Matsui (2003) Auditory morphing based on an elastic perceptual distance metric in an interference-free time-frequency representation. IEEE International Conference 1: I–I.

Lamm C, Decety J, Singer T (2011) Meta-analytic evidence for common and distinct neural networks associated with directly experienced pain and empathy for pain. Neuroimage 54:2492–2502.

LeDoux JE (2017) Semantics, surplus meaning, and the science of fear. Trends in cognitive sciences. 21: 303–306.

Menon V, Uddin LQ (2010) Saliency, switching, attention and control: a network model of insula function. Brain Struct. Funct. 214: 655–667.

Mesgarani N, Cheung C, Johnson K, Chang EF (2014) Phonetic feature encoding in human superior temporal gyrus. Science, 343:1006–1010.

Mesulam MM, Mufson EJ (1985) The insula of Reil in man and monkey: Architectonics, connectivity, and function. In: Peters A, Jones EG, editors. Cereb. Cortex, Vol. 4. Association and auditory cortices. New York: Plenum. 179–226.

Morningstar M, Nelson EE, & Dirks MA (2018) Maturation of vocal emotion recognition: Insights from the developmental and neuroimaging literature. Neuroscience & Biobehavioral Reviews. 90: 221–230.

Morris JS, Frith CD, Perrett DI, Rowland D, Young AW, Calder AJ, & Dolan RJ (1996) A differential neural response in the human amygdala to fearful and happy facial expressions. Nature. 383: 812–815.

Morris JS, Öhman A, Dolan RJ (1998) Conscious and unconscious emotional learning in the human amygdala. Nature. 393: 467–470.

Morris JS, Scott SK, Dolan RJ (1999) Saying it with feeling: neural responses to emotional vocalizations. Neuropsychologia. 37: 1155–1163.

Mufson EJ, Mesulam MM, Pandya DN (1981) Insular interconnections with the amygdala in the rhesus monkey. Neuroscience. 6: 1231–1248.

Mukamel R, Gelbard H, Arieli A, Hasson U, Fried I, & Malach R (2005) Coupling between neuronal firing, field potentials, and FMRI in human auditory cortex. Science. 309: 951–954.

Nir Y, Fisch L, Mukamel R, Gelbard-Sagiv H, Arieli A, Fried I, & Malach R (2007) Coupling between neuronal firing rate, gamma LFP, and BOLD fMRI is related to interneuronal correlations. Current Biology. 17: 1275–1285.

Ostrowsky K, Magnin M, Ryvlin P, Isnard J, Guenot M, Mauguiere F (2002) Representation of pain and somatic sensation in the human insula: a study of responses to direct electrical cortical stimulation. Cerebral Cortex. 12:376–385.

Phan KL, Wager T, Taylor SF, Liberzon I (2002) Functional neuroanatomy of emotion: a meta-analysis of emotion activation studies in PET and fMRI. Neuroimage. 16: 331–348.

Phillips ML, Young AW, Scott S, Calder AJ, Andrew C, Giampietro V, … & Gray JA (1998) Neural responses to facial and vocal expressions of fear and disgust. Proc. R. Soc. Lond. B. Biol. Sci. 265: 1809–1817.

Rauschecker JP, Scott SK (2009) Maps and streams in the auditory cortex: nonhuman primates illuminate human speech processing. Nat. Neurosci. 12: 718–724.

Remedios R, Logothetis NK, Kayser C (2009) An auditory region in the primate insular cortex responding preferentially to vocal communication sounds. J. Neurosci. 29: 1034–1045.

Sander K, Scheich H (2001) Auditory perception of laughing and crying activates human amygdala regardless of attentional state. Brain Res. Cogn. Brain Res. 12: 181–198.

Schirmer A, & Kotz, SA (2006) Beyond the right hemisphere: brain mechanisms mediating vocal emotional processing. Trends in cognitive sciences. 10: 24–30.

Schweinhardt P, Glynn C, Brooks J, McQuay H, Jack T, Chessell I, Bountra C, Tracey I (2006) An fMRI study of cerebral processing of brush-evoked allodynia in neuropathic pain patients. Neuroimage. 32: 256–265.

Shelley BP, Trimble MR (2004) The insular lobe of Reil—its anatamico-functional, behavioural and neuropsychiatric attributes in humans—a review. World J Biol Psychiatry. 5:176–200.

Singer T, Seymour B, O’doherty J, Kaube H, Dolan RJ, Frith CD (2004) Empathy for pain involves the affective but not sensory components of pain. Science. 303:1157–1162.

Sridharan D, Levitin DJ, Menon V (2008) A critical role for the right fronto-insular cortex in switching between central-executive and default-mode networks. Proc. Natl. Acad. Sci. U.S.A. 105: 12569–12574.

Wager TD, Atlas LY, Lindquist MA, Roy M, Woo CW, Kross E (2013) An fMRI-based neurologic signature of physical pain. N Engl J Med. 368:1388–1397.

Wang S, Tudusciuc O, Mamelak AN, Ross IB, Adolphs R, Rutishauser U (2014) Neurons in the human amygdala selective for perceived emotion. Proc. Natl. Acad. Sci. U.S.A. 111: 3110–3119.

Wicker B, Keysers C, Plailly J, Royet JP, Gallese V, Rizzolatti G (2003) Both of us disgusted in my insula: the common neural basis of seeing and feeling disgust. Neuron. 40:655–664.

Wildgruber D, Ackermann H, Kreifelts B, & Ethofer T (2006) Cerebral processing of linguistic and emotional prosody: fMRI studies. Progress in brain research. 156: 249–268.

Yovel G, & Belin P (2013) A unified coding strategy for processing faces and voices. Trends in cognitive sciences. 17: 263–271.

Zaki J, Davis JI, Ochsner KN (2012) Overlapping activity in anterior insula during interoception and emotional experience. Neuroimage. 62: 493–499.

Zhang Y, Ding Y, Huang J, Zhou W, Ling Z, Hong B, Wang X (2021) Hierarchical cortical networks of “voice patches” for processing voices in human brain. Proc. Natl. Acad. Sci. U.S.A. 118(52).

Zhang Y, Zhou W, Wang S, Zhou Q, Wang H, Zhang B, … Wang X (2019) The roles of subdivisions of human insula in emotion perception and auditory processing. Cerebral Cortex. 29: 517–528.

