## Supplemental Material for "Neural correlates of perceived emotions in human insula and amygdala"

Figure S1

Analysis pipeline for Emotion Recognition Experiment

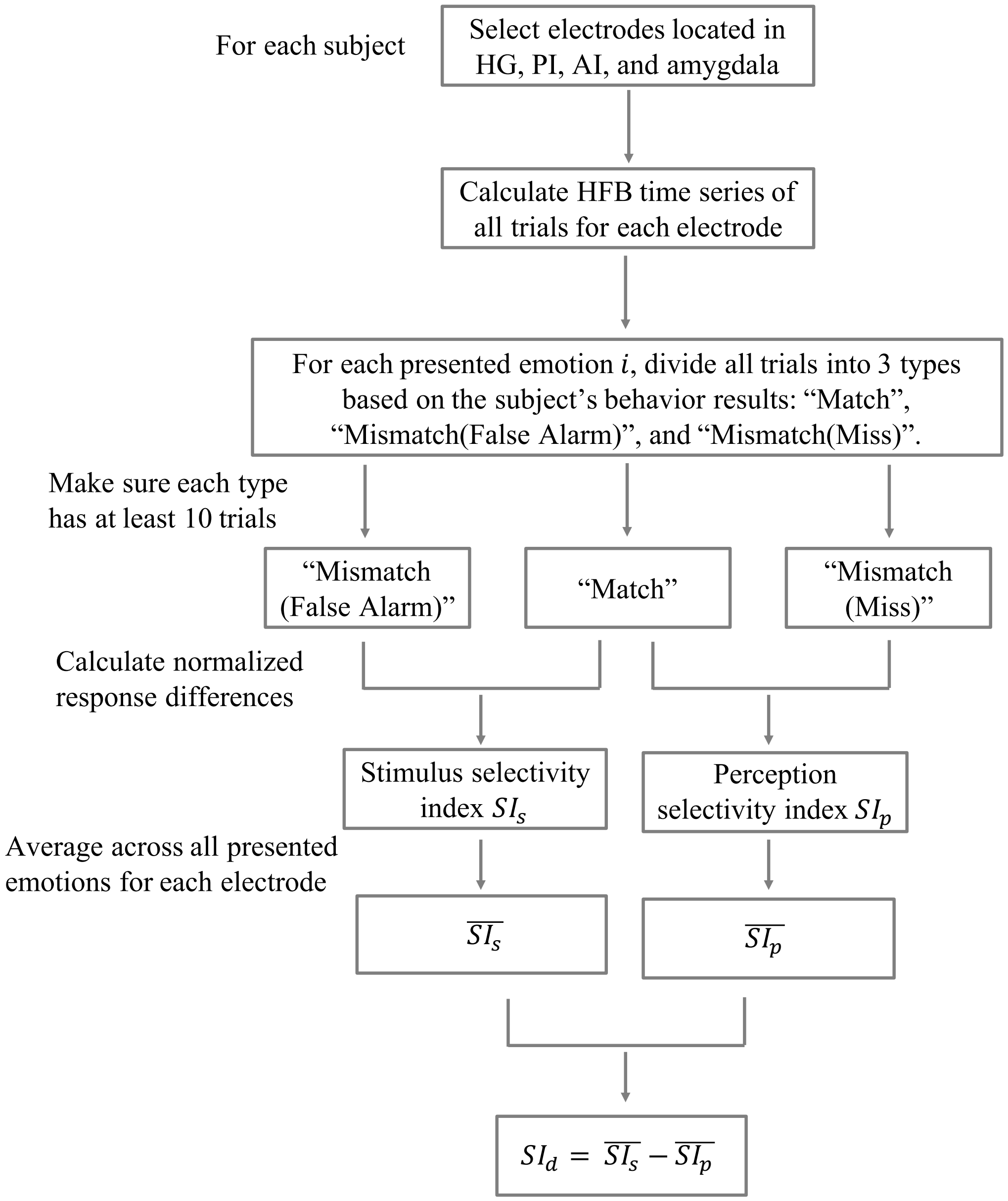


Each electrode has a $SI_d$ value to imply its preference. $SI_d > 0$ ($\overline{SI_s} > \overline{SI_p}$) indicates the electrode is selective to the acoustic properties of the stimuli, while $SI_d < 0$ ($\overline{SI_s} < \overline{SI_p}$) indicates the electrode is selective to subjective perceptual judgments of the perceivers.

Details of emotion recognition behavior results and the three types of trials from all subjects

"Match": Red

"Mismatch (Miss)": Green

"Mismatch (False Alarm)": Yellow

Numbers in the table: Trial number

Trial regroup criterion: Each type has at least 10 trials

**Subject 1:**

| | | Perceived Emotion | | | | | |
|---|---|---|---|---|---|---|---|
| Presented Emotion | | Anger | Disgust | Fear | Happy | Neutral | Sad |
| | Anger | 12 | 1 | 5 | 0 | 4 | 18 |
| | Disgust | 12 | 0 | 0 | 3 | 2 | 23 |
| | Fear | 15 | 0 | 0 | 0 | 5 | 20 |
| | Happy | 1 | 0 | 0 | 36 | 2 | 1 |
| | Neutral | 12 | 0 | 2 | 0 | 25 | 1 |
| | Sad | 1 | 0 | 0 | 0 | 1 | 38 |

Two conditions match the trial regroup criterion:

1: Presented emotion = Anger

| | | Perceived Emotion | | | | | |
|---|---|---|---|---|---|---|---|
| Presented Emotion | | Anger | Disgust | Fear | Happy | Neutral | Sad |
| | Anger | 12 | 1 | 5 | 0 | 4 | 18 |
| | Disgust | 12 | 0 | 0 | 3 | 2 | 23 |
| | Fear | 15 | 0 | 0 | 0 | 5 | 20 |
| | Happy | 1 | 0 | 0 | 36 | 2 | 1 |
| | Neutral | 12 | 0 | 2 | 0 | 25 | 1 |
| | Sad | 1 | 0 | 0 | 0 | 1 | 38 |

"Match" trial number = 12; "Mismatch (Miss)" trial number = 28; "Mismatch (False Alarm)" trial number = 41.

2: Presented emotion = Neutral

| | | Perceived Emotion | | | | | |
|---|---|---|---|---|---|---|---|
| Presented Emotion | | Anger | Disgust | Fear | Happy | Neutral | Sad |
| | Anger | 12 | 1 | 5 | 0 | 4 | 18 |
| | Disgust | 12 | 0 | 0 | 3 | 2 | 23 |
| | Fear | 15 | 0 | 0 | 0 | 5 | 20 |
| | Happy | 1 | 0 | 0 | 36 | 2 | 1 |
| | Neutral | 12 | 0 | 2 | 0 | 25 | 1 |
| | Sad | 1 | 0 | 0 | 0 | 1 | 38 |

"Match" trial number = 25; "Mismatch (Miss)" trial number = 15; "Mismatch (False Alarm)" trial number = 14.

**Subject 2:**

| | Perceived Emotion | | | | | | |
|---|---|---|---|---|---|---|---|
| Presented Emotion | | Anger | Disgust | Fear | Happy | Neutral | Sad |
| | Anger | 6 | 7 | 6 | 6 | 8 | 7 |
| | Disgust | 7 | 9 | 7 | 5 | 6 | 6 |
| | Fear | 3 | 5 | 7 | 10 | 4 | 11 |
| | Happy | 5 | 7 | 8 | 7 | 8 | 5 |
| | Neutral | 9 | 5 | 8 | 8 | 7 | 3 |
| | Sad | 8 | 7 | 7 | 5 | 6 | 7 |

No condition matches the trial regroup criterion.

**Subject 3:**

| | Perceived Emotion | | | | | | |
|---|---|---|---|---|---|---|---|
| Presented Emotion | | Anger | Disgust | Fear | Happy | Neutral | Sad |
| | Anger | 26 | 13 | 0 | 0 | 0 | 1 |
| | Disgust | 4 | 11 | 0 | 0 | 0 | 25 |
| | Fear | 2 | 0 | 38 | 0 | 0 | 0 |
| | Happy | 0 | 0 | 0 | 40 | 0 | 0 |
| | Neutral | 0 | 2 | 0 | 0 | 0 | 38 |
| | Sad | 1 | 1 | 6 | 1 | 31 | 0 |

One condition matches the trial regroup criterion.

1: Presented emotion = Disgust

| | Perceived Emotion | | | | | | |
|---|---|---|---|---|---|---|---|
| Presented Emotion | | Anger | Disgust | Fear | Happy | Neutral | Sad |
| | Anger | 26 | 13 | 0 | 0 | 0 | 1 |
| | Disgust | 4 | 11 | 0 | 0 | 0 | 25 |
| | Fear | 2 | 0 | 38 | 0 | 0 | 0 |
| | Happy | 0 | 0 | 0 | 40 | 0 | 0 |
| | Neutral | 0 | 2 | 0 | 0 | 0 | 38 |
| | Sad | 1 | 1 | 6 | 1 | 31 | 0 |

“Match” trial number = 11; “Mismatch (Miss)” trial number = 29; “Mismatch (False Alarm)” trial number = 16.

**Subject 4:**

| | Perceived Emotion | | | | | | |
|---|---|---|---|---|---|---|---|
| Presented Emotion | | Anger | Disgust | Fear | Happy | Neutral | Sad |
| | Anger | 8 | 5 | 27 | 0 | 0 | 0 |
| | Disgust | 2 | 17 | 4 | 0 | 0 | 17 |
| | Fear | 0 | 0 | 37 | 0 | 1 | 2 |
| | Happy | 0 | 0 | 0 | 40 | 0 | 0 |
| | Neutral | 3 | 27 | 4 | 0 | 0 | 6 |
| | Sad | 0 | 1 | 0 | 0 | 39 | 0 |

One condition matches the trial regroup criterion.

1: Presented emotion = Disgust

| | Perceived Emotion | | | | | | |
|---|---|---|---|---|---|---|---|
| | | Anger | Disgust | Fear | Happy | Neutral | Sad |
| Presented Emotion | Anger | 8 | 5 | 27 | 0 | 0 | 0 |
| | Disgust | 2 | 17 | 4 | 0 | 0 | 17 |
| | Fear | 0 | 0 | 37 | 0 | 1 | 2 |
| | Happy | 0 | 0 | 0 | 40 | 0 | 0 |
| | Neutral | 3 | 27 | 4 | 0 | 0 | 6 |
| | Sad | 0 | 1 | 0 | 0 | 39 | 0 |

“Match” trial number = 17; “Mismatch (Miss)” trial number = 23; “Mismatch (False Alarm)” trial number = 33.

**Subject 5:**

| | Perceived Emotion | | | | | | |
|---|---|---|---|---|---|---|---|
| | | Anger | Disgust | Fear | Happy | Neutral | Sad |
| Presented Emotion | Anger | 35 | 0 | 4 | 0 | 1 | 0 |
| | Disgust | 0 | 40 | 0 | 0 | 0 | 0 |
| | Fear | 0 | 0 | 39 | 0 | 1 | 0 |
| | Happy | 0 | 0 | 0 | 40 | 0 | 0 |
| | Neutral | 0 | 0 | 0 | 0 | 40 | 0 |
| | Sad | 0 | 0 | 0 | 0 | 0 | 40 |

No condition matches the trial regroup criterion.

**Subject 6:**

| | Perceived Emotion | | | | | | |
|---|---|---|---|---|---|---|---|
| | | Anger | Disgust | Fear | Happy | Neutral | Sad |
| Presented Emotion | Anger | 40 | 0 | 0 | 0 | 0 | 0 |
| | Disgust | 1 | 0 | 0 | 0 | 34 | 5 |
| | Fear | 0 | 0 | 40 | 0 | 0 | 0 |
| | Happy | 0 | 0 | 0 | 40 | 0 | 0 |
| | Neutral | 0 | 0 | 0 | 0 | 12 | 28 |
| | Sad | 0 | 39 | 0 | 1 | 0 | 0 |

One condition matches the trial regroup criterion.

1: Presented emotion = Neutral

| | Perceived Emotion | | | | | | |
|---|---|---|---|---|---|---|---|
| | | Anger | Disgust | Fear | Happy | Neutral | Sad |
| Presented Emotion | Anger | 40 | 0 | 0 | 0 | 0 | 0 |
| | Disgust | 1 | 0 | 0 | 0 | 34 | 5 |
| | Fear | 0 | 0 | 40 | 0 | 0 | 0 |
| | Happy | 0 | 0 | 0 | 40 | 0 | 0 |
| | Neutral | 0 | 0 | 0 | 0 | 12 | 28 |
| | Sad | 0 | 39 | 0 | 1 | 0 | 0 |

"Match" trial number = 12; "Mismatch (Miss)" trial number = 28; "Mismatch (False Alarm)" trial number = 34.

**Subject 7:**

| | | Perceived Emotion | | | | | |
|---|---|---|---|---|---|---|---|
| Presented Emotion | | Anger | Disgust | Fear | Happy | Neutral | Sad |
| | Anger | 40 | 0 | 0 | 0 | 0 | 0 |
| | Disgust | 0 | 40 | 0 | 0 | 0 | 0 |
| | Fear | 0 | 0 | 39 | 0 | 0 | 1 |
| | Happy | 0 | 0 | 0 | 40 | 0 | 0 |
| | Neutral | 0 | 0 | 0 | 0 | 0 | 40 |
| | Sad | 0 | 0 | 0 | 4 | 36 | 0 |

No condition matches the trial regroup criterion.

**Subject 8:**

| | | Perceived Emotion | | | | | |
|---|---|---|---|---|---|---|---|
| Presented Emotion | | Anger | Disgust | Fear | Happy | Neutral | Sad |
| | Anger | 10 | 4 | 1 | 5 | 6 | 14 |
| | Disgust | 2 | 20 | 17 | 0 | 0 | 1 |
| | Fear | 4 | 23 | 4 | 1 | 6 | 2 |
| | Happy | 11 | 6 | 3 | 4 | 8 | 8 |
| | Neutral | 19 | 1 | 0 | 4 | 5 | 11 |
| | Sad | 5 | 21 | 9 | 0 | 4 | 1 |

Two conditions match the trial regroup criterion:

1: Presented emotion = Anger

| | | Perceived Emotion | | | | | |
|---|---|---|---|---|---|---|---|
| Presented Emotion | | Anger | Disgust | Fear | Happy | Neutral | Sad |
| | Anger | 10 | 4 | 1 | 5 | 6 | 14 |
| | Disgust | 2 | 20 | 17 | 0 | 0 | 1 |
| | Fear | 4 | 23 | 4 | 1 | 6 | 2 |
| | Happy | 11 | 6 | 3 | 4 | 8 | 8 |
| | Neutral | 19 | 1 | 0 | 4 | 5 | 11 |
| | Sad | 5 | 21 | 9 | 0 | 4 | 1 |

"Match" trial number = 10; "Mismatch (Miss)" trial number = 30; "Mismatch (False Alarm)" trial number = 41.

2: Presented emotion = Disgust

| | | Perceived Emotion | | | | | |
|---|---|---|---|---|---|---|---|
| Presented Emotion | | Anger | Disgust | Fear | Happy | Neutral | Sad |
| | Anger | 10 | 4 | 1 | 5 | 6 | 14 |
| | Disgust | 2 | 20 | 17 | 0 | 0 | 1 |
| | Fear | 4 | 23 | 4 | 1 | 6 | 2 |

| | | | | | | | |
|---|---|---|---|---|---|---|---|
| | Happy | 11 | 6 | 3 | 4 | 8 | 8 |
| | Neutral | 19 | 1 | 0 | 4 | 5 | 11 |
| | Sad | 5 | 21 | 9 | 0 | 4 | 1 |

"Match" trial number = 20; "Mismatch (Miss)" trial number = 20; "Mismatch (False Alarm)" trial number = 55.

**Subject 9:**

| | Perceived Emotion | | | | | | |
|---|---|---|---|---|---|---|---|
| Presented Emotion | | Anger | Disgust | Fear | Happy | Neutral | Sad |
| | Anger | 39 | 1 | 0 | 0 | 0 | 0 |
| | Disgust | 0 | 18 | 1 | 0 | 0 | 21 |
| | Fear | 0 | 2 | 38 | 0 | 0 | 0 |
| | Happy | 0 | 1 | 0 | 39 | 0 | 0 |
| | Neutral | 0 | 40 | 0 | 0 | 0 | 0 |
| | Sad | 0 | 0 | 1 | 0 | 39 | 0 |

One condition matches the trial regroup criterion:

1: Presented emotion = Disgust

| | Perceived Emotion | | | | | | |
|---|---|---|---|---|---|---|---|
| Presented Emotion | | Anger | Disgust | Fear | Happy | Neutral | Sad |
| | Anger | 39 | 1 | 0 | 0 | 0 | 0 |
| | Disgust | 0 | 18 | 1 | 0 | 0 | 21 |
| | Fear | 0 | 2 | 38 | 0 | 0 | 0 |
| | Happy | 0 | 1 | 0 | 39 | 0 | 0 |
| | Neutral | 0 | 40 | 0 | 0 | 0 | 0 |
| | Sad | 0 | 0 | 1 | 0 | 39 | 0 |

"Match" trial number = 18; "Mismatch (Miss)" trial number = 22; "Mismatch (False Alarm)" trial number = 44.

**Subject 10:**

| | Perceived Emotion | | | | | | |
|---|---|---|---|---|---|---|---|
| Presented Emotion | | Anger | Disgust | Fear | Happy | Neutral | Sad |
| | Anger | 18 | 0 | 17 | 2 | 3 | 0 |
| | Disgust | 10 | 25 | 2 | 0 | 0 | 3 |
| | Fear | 0 | 4 | 32 | 0 | 4 | 0 |
| | Happy | 0 | 0 | 0 | 39 | 1 | 0 |
| | Neutral | 2 | 0 | 0 | 0 | 1 | 37 |
| | Sad | 0 | 0 | 0 | 2 | 38 | 0 |

One condition matches the trial regroup criterion:

1: Presented emotion = Anger

| | Perceived Emotion | | | | | | |
|---|---|---|---|---|---|---|---|
| | | Anger | Disgust | Fear | Happy | Neutral | Sad |
| | Anger | 18 | 0 | 17 | 2 | 3 | 0 |

| | | | | | | | |
|---|---|---|---|---|---|---|---|
| Presented Emotion | Disgust | 10 | 25 | 2 | 0 | 0 | 3 |
| | Fear | 0 | 4 | 32 | 0 | 4 | 0 |
| | Happy | 0 | 0 | 0 | 39 | 1 | 0 |
| | Neutral | 2 | 0 | 0 | 0 | 1 | 37 |
| | Sad | 0 | 0 | 0 | 2 | 38 | 0 |

"Match" trial number = 18; "Mismatch (Miss)" trial number = 22; "Mismatch (False Alarm)" trial number = 12.

**Subject 11:**

| | Perceived Emotion | | | | | | |
|---|---|---|---|---|---|---|---|
| | | Anger | Disgust | Fear | Happy | Neutral | Sad |
| Presented Emotion | Anger | 40 | 0 | 0 | 0 | 0 | 0 |
| | Disgust | 0 | 40 | 0 | 0 | 0 | 0 |
| | Fear | 1 | 0 | 39 | 0 | 0 | 0 |
| | Happy | 0 | 0 | 0 | 40 | 0 | 0 |
| | Neutral | 0 | 0 | 0 | 0 | 40 | 0 |
| | Sad | 0 | 0 | 0 | 0 | 0 | 40 |

No condition matches the trial regroup criterion.

**Subject 12:**

| | Perceived Emotion | | | | | | |
|---|---|---|---|---|---|---|---|
| | | Anger | Disgust | Fear | Happy | Neutral | Sad |
| Presented Emotion | Anger | 39 | 0 | 1 | 0 | 0 | 0 |
| | Disgust | 0 | 36 | 4 | 0 | 0 | 0 |
| | Fear | 0 | 0 | 40 | 0 | 0 | 0 |
| | Happy | 0 | 0 | 0 | 40 | 0 | 0 |
| | Neutral | 0 | 0 | 0 | 0 | 40 | 0 |
| | Sad | 0 | 0 | 0 | 0 | 0 | 40 |

No condition matches the trial regroup criterion.

**Subject 13:**

| | Perceived Emotion | | | | | | |
|---|---|---|---|---|---|---|---|
| | | Anger | Disgust | Fear | Happy | Neutral | Sad |
| Presented Emotion | Anger | 9 | 0 | 22 | 0 | 0 | 9 |
| | Disgust | 1 | 1 | 0 | 0 | 1 | 37 |
| | Fear | 0 | 0 | 29 | 0 | 0 | 11 |
| | Happy | 0 | 0 | 0 | 40 | 0 | 0 |
| | Neutral | 0 | 0 | 0 | 0 | 2 | 38 |
| | Sad | 3 | 0 | 0 | 3 | 34 | 0 |

One condition matches the trial regroup criterion:

1: Presented emotion = Fear

| | Perceived Emotion |
|---|---|

| | | Anger | Disgust | Fear | Happy | Neutral | Sad |
|---|---|---|---|---|---|---|---|
| Presented Emotion | Anger | 9 | 0 | 22 | 0 | 0 | 9 |
| | Disgust | 1 | 1 | 0 | 0 | 1 | 37 |
| | Fear | 0 | 0 | 29 | 0 | 0 | 11 |
| | Happy | 0 | 0 | 0 | 40 | 0 | 0 |
| | Neutral | 0 | 0 | 0 | 0 | 2 | 38 |
| | Sad | 3 | 0 | 0 | 3 | 34 | 0 |

"Match" trial number = 29; "Mismatch (Miss)" trial number = 11; "Mismatch (False Alarm)" trial number = 22.

**Subject 14:**

| | | Perceived Emotion | | | | | |
|---|---|---|---|---|---|---|---|
| | | Anger | Disgust | Fear | Happy | Neutral | Sad |
| Presented Emotion | Anger | 14 | 11 | 10 | 0 | 4 | 1 |
| | Disgust | 10 | 11 | 2 | 1 | 5 | 11 |
| | Fear | 2 | 12 | 19 | 0 | 2 | 5 |
| | Happy | 2 | 1 | 0 | 25 | 8 | 4 |
| | Neutral | 2 | 4 | 5 | 2 | 4 | 23 |
| | Sad | 1 | 3 | 1 | 0 | 35 | 0 |

Three conditions match the trial regroup criterion:

1: Presented emotion = Anger

| | | Perceived Emotion | | | | | |
|---|---|---|---|---|---|---|---|
| | | Anger | Disgust | Fear | Happy | Neutral | Sad |
| Presented Emotion | Anger | 14 | 11 | 10 | 0 | 4 | 1 |
| | Disgust | 10 | 11 | 2 | 1 | 5 | 11 |
| | Fear | 2 | 12 | 19 | 0 | 2 | 5 |
| | Happy | 2 | 1 | 0 | 25 | 8 | 4 |
| | Neutral | 2 | 4 | 5 | 2 | 4 | 23 |
| | Sad | 1 | 3 | 1 | 0 | 35 | 0 |

"Match" trial number = 14; "Mismatch (Miss)" trial number = 26; "Mismatch (False Alarm)" trial number = 17.

2: Presented emotion = Disgust

| | | Perceived Emotion | | | | | |
|---|---|---|---|---|---|---|---|
| | | Anger | Disgust | Fear | Happy | Neutral | Sad |
| Presented Emotion | Anger | 14 | 11 | 10 | 0 | 4 | 1 |
| | Disgust | 10 | 11 | 2 | 1 | 5 | 11 |
| | Fear | 2 | 12 | 19 | 0 | 2 | 5 |
| | Happy | 2 | 1 | 0 | 25 | 8 | 4 |
| | Neutral | 2 | 4 | 5 | 2 | 4 | 23 |
| | Sad | 1 | 3 | 1 | 0 | 35 | 0 |

"Match" trial number = 11; "Mismatch (Miss)" trial number = 29; "Mismatch (False Alarm)" trial number = 31.

3: Presented emotion = Fear

| | Perceived Emotion | | | | | | |
|---|---|---|---|---|---|---|---|
| | | Anger | Disgust | Fear | Happy | Neutral | Sad |
| Presented Emotion | Anger | 14 | 11 | 10 | 0 | 4 | 1 |
| | Disgust | 10 | 11 | 2 | 1 | 5 | 11 |
| | Fear | 2 | 12 | 19 | 0 | 2 | 5 |
| | Happy | 2 | 1 | 0 | 25 | 8 | 4 |
| | Neutral | 2 | 4 | 5 | 2 | 4 | 23 |
| | Sad | 1 | 3 | 1 | 0 | 35 | 0 |

"Match" trial number = 19; "Mismatch (Miss)" trial number = 21; "Mismatch (False Alarm)" trial number = 18.

**Subject 15:**

| | Perceived Emotion | | | | | | |
|---|---|---|---|---|---|---|---|
| | | Anger | Disgust | Fear | Happy | Neutral | Sad |
| Presented Emotion | Anger | 33 | 1 | 4 | 0 | 0 | 2 |
| | Disgust | 0 | 36 | 0 | 0 | 0 | 4 |
| | Fear | 0 | 0 | 39 | 0 | 0 | 1 |
| | Happy | 0 | 0 | 0 | 40 | 0 | 0 |
| | Neutral | 0 | 3 | 6 | 0 | 0 | 31 |
| | Sad | 0 | 0 | 1 | 0 | 39 | 0 |

No condition matches the trial regroup criterion.

**Subject 16:**

| | Perceived Emotion | | | | | | |
|---|---|---|---|---|---|---|---|
| | | Anger | Disgust | Fear | Happy | Neutral | Sad |
| Presented Emotion | Anger | 39 | 0 | 1 | 0 | 0 | 0 |
| | Disgust | 0 | 0 | 0 | 0 | 2 | 38 |
| | Fear | 0 | 0 | 40 | 0 | 0 | 0 |
| | Happy | 0 | 0 | 0 | 40 | 0 | 0 |
| | Neutral | 0 | 40 | 0 | 0 | 0 | 0 |
| | Sad | 0 | 0 | 0 | 0 | 20 | 20 |

One condition matches the trial regroup criterion:

1: Presented emotion = Sad

| | Perceived Emotion | | | | | | |
|---|---|---|---|---|---|---|---|
| | | Anger | Disgust | Fear | Happy | Neutral | Sad |
| Presented Emotion | Anger | 39 | 0 | 1 | 0 | 0 | 0 |
| | Disgust | 0 | 0 | 0 | 0 | 2 | 38 |
| | Fear | 0 | 0 | 40 | 0 | 0 | 0 |
| | Happy | 0 | 0 | 0 | 40 | 0 | 0 |
| | Neutral | 0 | 40 | 0 | 0 | 0 | 0 |
| | Sad | 0 | 0 | 0 | 0 | 20 | 20 |

"Match" trial number = 20; "Mismatch (Miss)" trial number = 20; "Mismatch (False Alarm)" trial number = 38.

**Subject 17:**

| | | Perceived Emotion | | | | | |
|---|---|---|---|---|---|---|---|
| Presented Emotion | | Anger | Disgust | Fear | Happy | Neutral | Sad |
| | Anger | 20 | 0 | 0 | 0 | 0 | 20 |
| | Disgust | 0 | 0 | 0 | 0 | 0 | 40 |
| | Fear | 0 | 0 | 23 | 0 | 0 | 17 |
| | Happy | 0 | 0 | 0 | 38 | 0 | 2 |
| | Neutral | 0 | 0 | 0 | 0 | 0 | 40 |
| | Sad | 0 | 0 | 0 | 3 | 17 | 20 |

One condition matches the trial regroup criterion:

1: Presented emotion = Sad

| | | Perceived Emotion | | | | | |
|---|---|---|---|---|---|---|---|
| Presented Emotion | | Anger | Disgust | Fear | Happy | Neutral | Sad |
| | Anger | 20 | 0 | 0 | 0 | 0 | 20 |
| | Disgust | 0 | 0 | 0 | 0 | 0 | 40 |
| | Fear | 0 | 0 | 23 | 0 | 0 | 17 |
| | Happy | 0 | 0 | 0 | 38 | 0 | 2 |
| | Neutral | 0 | 0 | 0 | 0 | 0 | 40 |
| | Sad | 0 | 0 | 0 | 3 | 17 | 20 |

"Match" trial number = 20; "Mismatch (Miss)" trial number = 20; "Mismatch (False Alarm)" trial number = 119.

**Subject 18:**

| | | Perceived Emotion | | | | | |
|---|---|---|---|---|---|---|---|
| Presented Emotion | | Anger | Disgust | Fear | Happy | Neutral | Sad |
| | Anger | 32 | 1 | 2 | 0 | 0 | 5 |
| | Disgust | 1 | 25 | 2 | 0 | 0 | 12 |
| | Fear | 0 | 5 | 32 | 0 | 1 | 2 |
| | Happy | 0 | 0 | 0 | 38 | 2 | 0 |
| | Neutral | 0 | 2 | 1 | 0 | 0 | 37 |
| | Sad | 0 | 1 | 1 | 1 | 37 | 0 |

No condition matches the trial regroup criterion.

**Subject 19:**

| | | Perceived Emotion | | | | | |
|---|---|---|---|---|---|---|---|
| Presented Emotion | | Anger | Disgust | Fear | Happy | Neutral | Sad |
| | Anger | 31 | 2 | 5 | 1 | 0 | 1 |
| | Disgust | 0 | 27 | 10 | 0 | 0 | 3 |
| | Fear | 20 | 0 | 17 | 0 | 0 | 3 |
| | Happy | 0 | 0 | 0 | 40 | 0 | 0 |
| | Neutral | 0 | 31 | 4 | 0 | 0 | 5 |
| | Sad | 0 | 1 | 8 | 10 | 18 | 3 |

Two conditions match the trial regroup criterion:

1: Presented emotion = Disgust

| | Perceived Emotion | | | | | | |
|---|---|---|---|---|---|---|---|
| | | Anger | Disgust | Fear | Happy | Neutral | Sad |
| Presented Emotion | Anger | 31 | 2 | 5 | 1 | 0 | 1 |
| | Disgust | 0 | 27 | 10 | 0 | 0 | 3 |
| | Fear | 20 | 0 | 17 | 0 | 0 | 3 |
| | Happy | 0 | 0 | 0 | 40 | 0 | 0 |
| | Neutral | 0 | 31 | 4 | 0 | 0 | 5 |
| | Sad | 0 | 1 | 8 | 10 | 18 | 3 |

"Match" trial number = 27; "Mismatch (Miss)" trial number = 13; "Mismatch (False Alarm)" trial number = 34.

2: Presented emotion = Fear

| | Perceived Emotion | | | | | | |
|---|---|---|---|---|---|---|---|
| | | Anger | Disgust | Fear | Happy | Neutral | Sad |
| Presented Emotion | Anger | 31 | 2 | 5 | 1 | 0 | 1 |
| | Disgust | 0 | 27 | 10 | 0 | 0 | 3 |
| | Fear | 20 | 0 | 17 | 0 | 0 | 3 |
| | Happy | 0 | 0 | 0 | 40 | 0 | 0 |
| | Neutral | 0 | 31 | 4 | 0 | 0 | 5 |
| | Sad | 0 | 1 | 8 | 10 | 18 | 3 |

"Match" trial number = 17; "Mismatch (Miss)" trial number = 23; "Mismatch (False Alarm)" trial number = 27.

**Subject 20:**

| | Perceived Emotion | | | | | | |
|---|---|---|---|---|---|---|---|
| | | Anger | Disgust | Fear | Happy | Neutral | Sad |
| Presented Emotion | Anger | 16 | 6 | 6 | 0 | 10 | 2 |
| | Disgust | 1 | 31 | 2 | 2 | 2 | 2 |
| | Fear | 2 | 4 | 25 | 1 | 2 | 6 |
| | Happy | 2 | 0 | 0 | 37 | 1 | 0 |
| | Neutral | 4 | 6 | 1 | 2 | 10 | 17 |
| | Sad | 30 | 2 | 2 | 3 | 2 | 1 |

Three conditions match the trial regroup criterion:

1: Presented emotion = Anger

| | Perceived Emotion | | | | | | |
|---|---|---|---|---|---|---|---|
| | | Anger | Disgust | Fear | Happy | Neutral | Sad |
| Presented Emotion | Anger | 16 | 6 | 6 | 0 | 10 | 2 |
| | Disgust | 1 | 31 | 2 | 2 | 2 | 2 |
| | Fear | 2 | 4 | 25 | 1 | 2 | 6 |
| | Happy | 2 | 0 | 0 | 37 | 1 | 0 |
| | Neutral | 4 | 6 | 1 | 2 | 10 | 17 |
| | Sad | 30 | 2 | 2 | 3 | 2 | 1 |

“Match” trial number = 16; “Mismatch (Miss)” trial number = 24; “Mismatch (False Alarm)” trial number = 39.

2: Presented emotion = Fear

| | | Perceived Emotion | | | | | |
|---|---|---|---|---|---|---|---|
| Presented Emotion | | Anger | Disgust | Fear | Happy | Neutral | Sad |
| | Anger | 16 | 6 | 6 | 0 | 10 | 2 |
| | Disgust | 1 | 31 | 2 | 2 | 2 | 2 |
| | Fear | 2 | 4 | 25 | 1 | 2 | 6 |
| | Happy | 2 | 0 | 0 | 37 | 1 | 0 |
| | Neutral | 4 | 6 | 1 | 2 | 10 | 17 |
| | Sad | 30 | 2 | 2 | 3 | 2 | 1 |

“Match” trial number = 25; “Mismatch (Miss)” trial number = 15; “Mismatch (False Alarm)” trial number = 11.

3: Presented emotion = Neutral

| | | Perceived Emotion | | | | | |
|---|---|---|---|---|---|---|---|
| Presented Emotion | | Anger | Disgust | Fear | Happy | Neutral | Sad |
| | Anger | 16 | 6 | 6 | 0 | 10 | 2 |
| | Disgust | 1 | 31 | 2 | 2 | 2 | 2 |
| | Fear | 2 | 4 | 25 | 1 | 2 | 6 |
| | Happy | 2 | 0 | 0 | 37 | 1 | 0 |
| | Neutral | 4 | 6 | 1 | 2 | 10 | 17 |
| | Sad | 30 | 2 | 2 | 3 | 2 | 1 |

“Match” trial number = 10; “Mismatch (Miss)” trial number = 30; “Mismatch (False Alarm)” trial number = 17.

**Subject 21:**

| | | Perceived Emotion | | | | | |
|---|---|---|---|---|---|---|---|
| Presented Emotion | | Anger | Disgust | Fear | Happy | Neutral | Sad |
| | Anger | 5 | 6 | 21 | 0 | 7 | 1 |
| | Disgust | 2 | 3 | 7 | 0 | 2 | 26 |
| | Fear | 2 | 20 | 13 | 0 | 4 | 1 |
| | Happy | 0 | 0 | 1 | 39 | 0 | 0 |
| | Neutral | 1 | 5 | 28 | 0 | 4 | 2 |
| | Sad | 3 | 1 | 4 | 0 | 28 | 4 |

One condition matches the trial regroup criterion:

1: Presented emotion = Fear

| | | Perceived Emotion | | | | | |
|---|---|---|---|---|---|---|---|
| Presented Emotion | | Anger | Disgust | Fear | Happy | Neutral | Sad |
| | Anger | 5 | 6 | 21 | 0 | 7 | 1 |
| | Disgust | 2 | 3 | 7 | 0 | 2 | 26 |
| | Fear | 2 | 20 | 13 | 0 | 4 | 1 |
| | Happy | 0 | 0 | 1 | 39 | 0 | 0 |

| | | Anger | Disgust | Fear | Happy | Neutral | Sad |
|---|---|---|---|---|---|---|---|
| | Neutral | 1 | 5 | 28 | 0 | 4 | 2 |
| | Sad | 3 | 1 | 4 | 0 | 28 | 4 |

"Match" trial number = 13; "Mismatch (Miss)" trial number = 27; "Mismatch (False Alarm)" trial number = 61.

**Subject 22:**

| | Perceived Emotion | | | | | | |
|---|---|---|---|---|---|---|---|
| Presented Emotion | | Anger | Disgust | Fear | Happy | Neutral | Sad |
| | Anger | 36 | 0 | 1 | 0 | 0 | 3 |
| | Disgust | 0 | 37 | 2 | 0 | 0 | 1 |
| | Fear | 1 | 1 | 37 | 0 | 0 | 1 |
| | Happy | 0 | 0 | 0 | 39 | 1 | 0 |
| | Neutral | 0 | 1 | 0 | 0 | 0 | 39 |
| | Sad | 0 | 0 | 0 | 0 | 40 | 0 |

No condition matches the trial regroup criterion.

**Subject 23:**

| | Perceived Emotion | | | | | | |
|---|---|---|---|---|---|---|---|
| Presented Emotion | | Anger | Disgust | Fear | Happy | Neutral | Sad |
| | Anger | 1 | 1 | 35 | 0 | 0 | 3 |
| | Disgust | 2 | 26 | 0 | 0 | 1 | 11 |
| | Fear | 11 | 3 | 24 | 0 | 0 | 2 |
| | Happy | 1 | 1 | 0 | 36 | 2 | 0 |
| | Neutral | 6 | 29 | 1 | 0 | 0 | 4 |
| | Sad | 7 | 7 | 0 | 5 | 16 | 5 |

Two conditions match the trial regroup criterion:

1: Presented emotion = Disgust

| | Perceived Emotion | | | | | | |
|---|---|---|---|---|---|---|---|
| Presented Emotion | | Anger | Disgust | Fear | Happy | Neutral | Sad |
| | Anger | 1 | 1 | 35 | 0 | 0 | 3 |
| | Disgust | 2 | 26 | 0 | 0 | 1 | 11 |
| | Fear | 11 | 3 | 24 | 0 | 0 | 2 |
| | Happy | 1 | 1 | 0 | 36 | 2 | 0 |
| | Neutral | 6 | 29 | 1 | 0 | 0 | 4 |
| | Sad | 7 | 7 | 0 | 5 | 16 | 5 |

"Match" trial number = 26; "Mismatch (Miss)" trial number = 14; "Mismatch (False Alarm)" trial number = 41.

2: Presented emotion = Fear

| | Perceived Emotion | | | | | | |
|---|---|---|---|---|---|---|---|
| | | Anger | Disgust | Fear | Happy | Neutral | Sad |
| | Anger | 1 | 1 | 35 | 0 | 0 | 3 |

| | | | | | | | |
|---|---|---|---|---|---|---|---|
| Presented Emotion | Disgust | 2 | 26 | 0 | 0 | 1 | 11 |
| | Fear | 11 | 3 | 24 | 0 | 0 | 2 |
| | Happy | 1 | 1 | 0 | 36 | 2 | 0 |
| | Neutral | 6 | 29 | 1 | 0 | 0 | 4 |
| | Sad | 7 | 7 | 0 | 5 | 16 | 5 |

“Match” trial number = 24; “Mismatch (Miss)” trial number = 16; “Mismatch (False Alarm)” trial number = 36.

**Subject 24:**

| | | Perceived Emotion | | | | | |
|---|---|---|---|---|---|---|---|
| | | Anger | Disgust | Fear | Happy | Neutral | Sad |
| Presented Emotion | Anger | 34 | 0 | 0 | 0 | 2 | 4 |
| | Disgust | 0 | 31 | 9 | 0 | 0 | 0 |
| | Fear | 3 | 3 | 15 | 0 | 2 | 17 |
| | Happy | 0 | 0 | 0 | 39 | 1 | 0 |
| | Neutral | 0 | 4 | 34 | 0 | 2 | 0 |
| | Sad | 1 | 1 | 0 | 0 | 38 | 0 |

One condition matches the trial regroup criterion:

1: Presented emotion = Fear

| | | Perceived Emotion | | | | | |
|---|---|---|---|---|---|---|---|
| | | Anger | Disgust | Fear | Happy | Neutral | Sad |
| Presented Emotion | Anger | 34 | 0 | 0 | 0 | 2 | 4 |
| | Disgust | 0 | 31 | 9 | 0 | 0 | 0 |
| | Fear | 3 | 3 | 15 | 0 | 2 | 17 |
| | Happy | 0 | 0 | 0 | 39 | 1 | 0 |
| | Neutral | 0 | 4 | 34 | 0 | 2 | 0 |
| | Sad | 1 | 1 | 0 | 0 | 38 | 0 |

“Match” trial number = 15; “Mismatch (Miss)” trial number = 25; “Mismatch (False Alarm)” trial number = 43.
